# Human mutations in high-confidence Tourette disorder genes affect sensorimotor behavior, reward learning, and striatal dopamine in mice

**DOI:** 10.1101/2023.11.28.569034

**Authors:** Cara Nasello, Lauren A. Poppi, Junbing Wu, Tess F. Kowalski, Joshua K. Thackray, Riley Wang, Angelina Persaud, Mariam Mahboob, Sherry Lin, Rodna Spaseska, C.K. Johnson, Derek Gordon, Fadel Tissir, Gary A. Heiman, Jay A. Tischfield, Miriam Bocarsly, Max A. Tischfield

**Affiliations:** Department of Genetics and the Human Genetics Institute of New Jersey, Rutgers University, Piscataway, NJ 08854, USA; Department of Cell Biology and Neuroscience, Rutgers University, Piscataway, NJ 08854, USA; Child Health Institute of New Jersey, Robert Wood Johnson Medical School, New Brunswick, NJ 08901, USA; Department of Pharmacology, Physiology, and Neuroscience, Rutgers New Jersey Medical School and Rutgers Biomedical and Health Sciences, Newark, NJ 07103, USA; Deparment of Neurobiology, Harvard Medical School, Boston, MA 02115; College of Health and Life Sciences, Hamad Bin Khalifa University, Doha, Qatar; Université Catholique de Louvain, Institute of Neuroscience, Brussels, Belgium

## Abstract

Tourette disorder (TD) is poorly understood, despite affecting 1/160 children. A lack of animal models possessing construct, face, and predictive validity hinders progress in the field. We used CRISPR/Cas9 genome editing to generate mice with mutations orthologous to human *de novo* variants in two high-confidence Tourette genes, *CELSR3* and *WWC1*. Mice with human mutations in *Celsr3* and *Wwc1* exhibit cognitive and/or sensorimotor behavioral phenotypes consistent with TD. Sensorimotor gating deficits, as measured by acoustic prepulse inhibition, occur in both male and female *Celsr3* TD models. *Wwc1* mice show reduced prepulse inhibition only in females. Repetitive motor behaviors, common to *Celsr3* mice and more pronounced in females, include vertical rearing and grooming. Sensorimotor gating deficits and rearing are attenuated by aripiprazole, a partial agonist at dopamine type II receptors. Unsupervised machine learning reveals numerous changes to spontaneous motor behavior and less predictable patterns of movement. Continuous fixed-ratio reinforcement shows *Celsr3* TD mice have enhanced motor responding and reward learning. Electrically evoked striatal dopamine release, tested in one model, is greater. Brain development is otherwise grossly normal without signs of striatal interneuron loss. Altogether, mice expressing human mutations in high-confidence TD genes exhibit face and predictive validity. Reduced prepulse inhibition and repetitive motor behaviors are core behavioral phenotypes and are responsive to aripiprazole. Enhanced reward learning and motor responding occurs alongside greater evoked dopamine release. Phenotypes can also vary by sex and show stronger affection in females, an unexpected finding considering males are more frequently affected in TD.

**Significance Statement:** We generated mouse models that express mutations in high-confidence genes linked to Tourette disorder (TD). These models show sensorimotor and cognitive behavioral phenotypes resembling TD-like behaviors. Sensorimotor gating deficits and repetitive motor behaviors are attenuated by drugs that act on dopamine. Reward learning and striatal dopamine is enhanced. Brain development is grossly normal, including cortical layering and patterning of major axon tracts. Further, no signs of striatal interneuron loss are detected. Interestingly, behavioral phenotypes in affected females can be more pronounced than in males, despite male sex bias in the diagnosis of TD. These novel mouse models with construct, face, and predictive validity provide a new resource to study neural substrates that cause tics and related behavioral phenotypes in TD.

## Introduction

Tourette disorder (TD) is characterized by tics; fast, recurrent movements and vocalizations that wax and wane over time. Tics are frequently precipitated by unpleasant sensations and urges, with the tic providing relief (1, 2). Chronic tic disorders (CTD), a broad category that includes TD, are estimated to affect ∼1/50 school aged children in the United States and China (3–5). Common comorbidities include obsessive-compulsive, attention-deficit hyperactivity, and autism spectrum disorders, in addition to mood, anxiety, and sleep disorders (6). The constellation of neurological and neuropsychiatric conditions associated with TD/CTD impact social wellbeing, scholastic performance, and overall quality of life, posing increased risk for substance abuse, experiencing violent assault, and/or suicide (7–10). Yet, treatments are limited and frequently cause unwanted side-effects.

Animal models are needed to help identify the underlying neuronal substrates, affected brain regions, and circuit mechanisms that cause tics and related behaviors in TD. However, ascribing changes in brain activity to behavioral phenotypes in the absence of a precise genetic change or neuronal lesion found in an affected human confounds our ability to confidently test these mechanisms. As such, a lack of animal models possessing construct validity for TD has hindered progress in the field. Most TD animal models reported to date either lack construct validity or attempt to model “TD-like” phenotypes in the absence of *bona fide* gene mutations that are associated with the human disorder. Developing animal models with true construct validity necessitates identifying genetic variants in high-confidence human risk genes.

The Tourette International Collaborative Genetics (TIC Genetics) study was established in 2011 to elucidate the genetic architecture of CTDs, including TD (11). Whole-exome sequencing of over 800 CTD/TD simplex/multiplex families identified multiple, individually distinct, rare *de novo* mutations in six genes from unrelated probands. These included likely gene-function disrupting single nucleotide variants, insertion-deletion mutations, and copy number variants (12, 13). Rare *de novo* variants are enriched in CTD/TD simplex trios and are estimated to produce greater than 10% of cases (12). Rare variants, as opposed to more common variants found in genome-wide association studies, have larger effect sizes and thus are strong candidates for animal models. Two putative TD genes that meet high-confidence criteria (false discovery rate [FDR] < 0.1), *CELSR3* and *WWC1* (also known as KIBRA), are intriguing candidates because of their roles in brain development, expression in cortico-striato-thalamo-cortical circuits, and/or synaptic functions (14–19).

We used CRISPR/Cas9 gene editing to engineer mice that express mutations identical to the *de novo* mutations found in *CELSR3* and *WWC1* from TD probands. These models possess construct validity, thereby constituting a significant advance in the field. Using traditional measures of mouse behavior, we characterize sensorimotor gating deficits, hyperactivity, and repetitive-like motor behaviors, which can show sex-dependent variable expressivity. Aripiprazole administration can attenuate sensorimotor gating deficits and repetitive-like rearing behavior. Using machine learning, we also see changes to the patterning and execution of spontaneous motor behavior. Instrumental learning paradigms suggest reward learning is enhanced in *Celsr3* models, and electrically evoked dopamine (DA) release is greater in the striatum. Brain development is otherwise grossly normal and we do not detect striatal interneuron loss. These novel genetic models with construct validity show how mutations in high-confidence TD genes impact mouse behavior, providing possible outcome phenotypes for medication trials.

## Results

### Generation of TD mouse models with true construct validity

CRISPR/Cas9 genome editing was used to introduce orthologous human mutations into coding exons for mouse *Celsr3* and *Wwc1* (12) (**Figs. 1A, D and S1A, B**). For *Celsr3*, we engineered a non-synonymous cysteine to tyrosine substitution within the second laminin G-like repeat, corresponding to human and mouse residues C1915Y (RefSeq: NP_001398.2) and C1906Y (RefSeq: NP_001346501.1), respectively. We also introduced a base-pair insertion corresponding to a frameshift mutation within the second laminin G-like repeat of human (coding 5706dupC, protein G1903Rfs*106) and mouse (coding 5679dupC, protein S1894Rfs*2). The human and mouse insertions cause stop-gain mutations that truncate the protein within the second laminin G-like repeat **(Fig. 1A and Fig. S1A)**. Heterozygous *Celsr3^C1906Y/+^*and *Celsr3^p.S1894Rfs*2/+^* mice were viable but homozygotes died at birth, consistent with *Celsr3* knockout mice and loss-of-function effects on the protein (14, 20). Celsr3 protein from whole brain lysates was reduced in *Celsr3^S1894Rfs*2/+^* mice but, interestingly, increased in *Celsr3^C1906Y/+^*mice, suggesting the latter mutation may exert dominant-negative effects. (**Fig. 1B, C**). For *Wwc1*, we introduced a non-synonymous tryptophan to cysteine substitution corresponding to human (RefSeq: NP_001155133.1) and mouse (RefSeq:NP_740749.1) residue W88C (**Fig. 1D and Fig. S1B**). Heterozygous *Wwc1^W88C/+^* mice were viable and fertile, and protein levels from whole brain lysates were normal (**Fig. 1E**).

**Figure 1:**
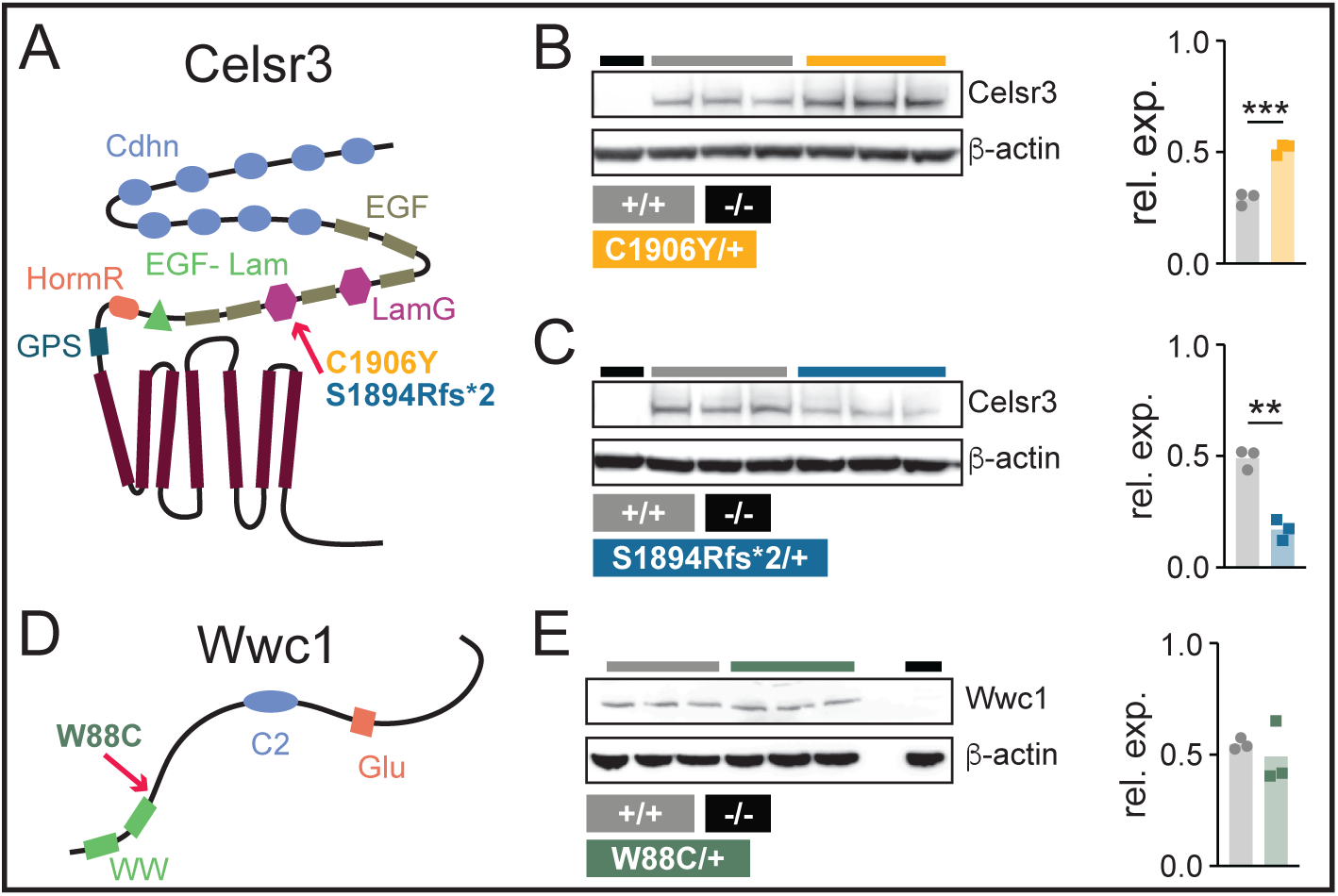
Celsr3 and Wwc1 protein domains, mutation sites, and relative levels. **(A)** Celsr3 protein schematic with domains and mutations denoted. Transmembrane (TM), G-protein signaling (GPS), hormone receptor (HormR), epidermal growth factor-like laminin (EGF-Lam), laminin G (LamG), epidermal growth factor (EGF), and cadherin (Cadhn) domains. **(B)** *Celsr3* KO (*Celsr3^-/-^*, lane 1), wild-types (*Celsr3^+/+^*, lanes 2-4), and mutants (*Celsr3^C1906Y/+^*, lanes 5-7). Celsr3 protein levels, relative to □-actin, are shown in bar plot (*p=0.0006*). **(C)** *Celsr3^-/-^* (lane 1), *Celsr3^+/+^* (lanes 2-4), *Celsr3^p.S1894Rfs*2/+^*(lanes 5-7). Celsr3 protein levels, relative to □-actin, are shown in bar plot (*p=0.001*). **(D)** Wwc1 protein schematic with WW and C2 domains, and a glutamic (Glu)-rich sequence. W88C substitution (green) is located after the second WW domain. **(E)** Wild-type (*Wwc1^+/+^*, lanes 1-3), mutant (*Wwc1^W88C/+^*, lanes 4-6), blank lane (X, lane 7), and homozygous null (*Wwc1^-/-^*, lane 8). Protein levels, relative to □-actin, are shown in bar plot (*p=0.534*). Unpaired t-test (B, C, E). *\*\*p<0.01, ***p<0.001*

### *Celsr3* and *Wwc1* mutant mice have sensorimotor gating deficits

Sensorimotor gating deficits, as measured by prepulse inhibition (PPI), are reported in humans with TD, and also in *Hdc* knockout mice which model a stop gain mutation found in a single TD family (21–24). We assessed whether *Celsr3* and *Wwc1* transgenic mice had deficits in PPI of the acoustic startle reflex (**Table S1**). Overall PPI was reduced in both *Celsr3^C1906Y/+^* females [F(1, 75)=16.95, *p=0.0005*] and males [F(1, 69)=13.46 *p<0.0001*] (**Fig. 2A**). Baseline acoustic startle responses were normal (**Fig. S2A**). PPI was also reduced in *Celsr3^S1894Rfs*2/+^* females [F(1, 108)=8.336 *p=0.0047*] and males [F(1, 75)=8.714, *p=0.0042*] (**Fig. 2B**). Baseline acoustic startle responses were normal in *Celsr3^S1894Rfs*2/+^* males, but decreased in females (*p=0.038)*. We did not detect an association between reduced startle and levels of inhibition (r=0.267 and *p=0.20*, Spearman’s correlation coefficient) (**Fig. S2B, C**). Acoustic PPI was also decreased in *Wwc1^W88C/+^* females [F(1, 105)=7.686, *p=0.0066*], who had normal baseline startle responses. *Wwc1^W88C/+^* males showed normal PPI and baseline startle responses [F(1,120)=0.8175, *p=0.37*] (**Figs. 2C, S2D**).

**Figure 2:**
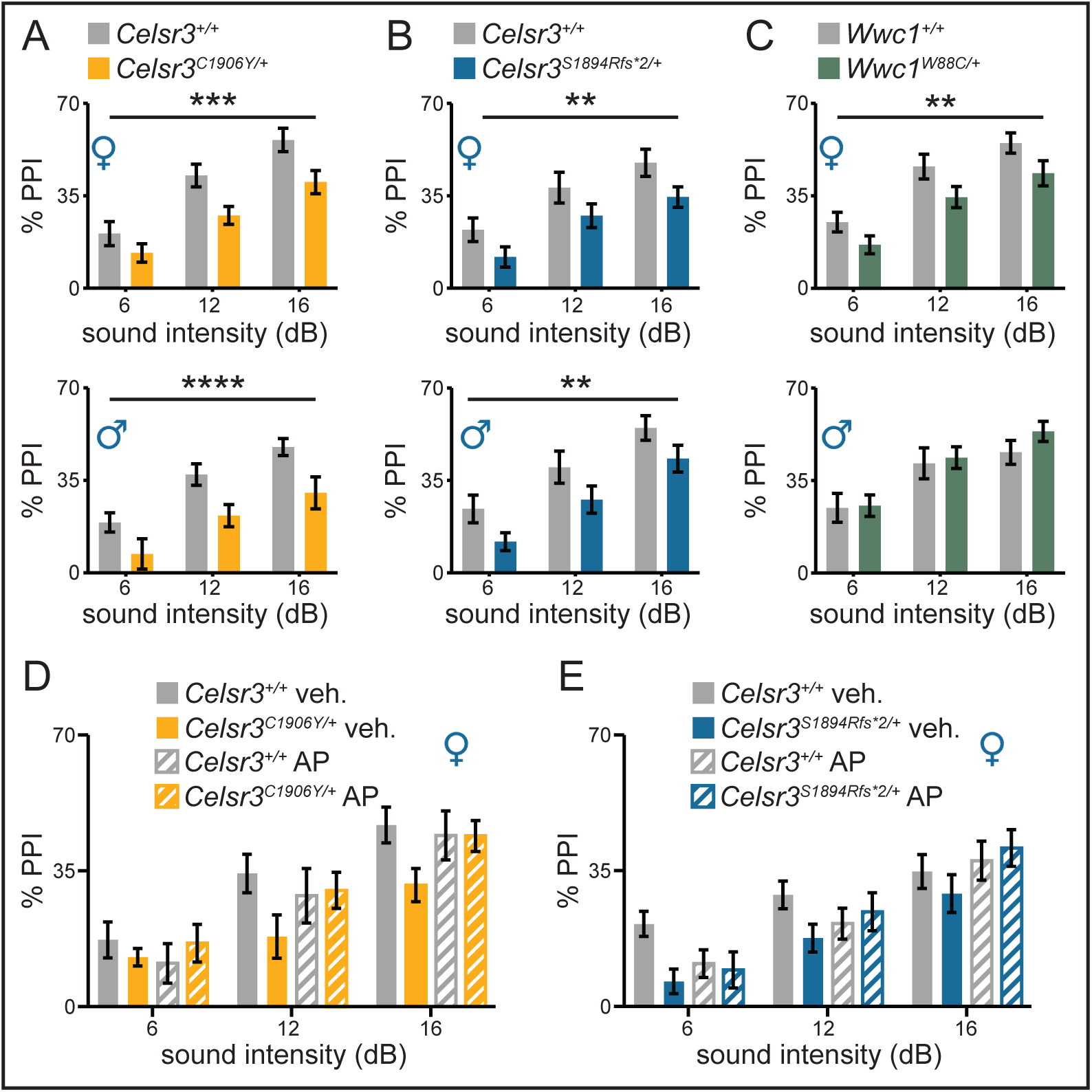
Celsr3 and Wwc1 mutant mice have reduced PPI. **(A)** Female *Celsr3^+/+^*/*Celsr3^C1906Y/+^* (n=14/11), *p=0.0005*; Male *Celsr3^+/+^*/*Celsr3^C1906Y/+^* (n=16/11), *p<0.0001*. **(B)** Female *Celsr3^+/+^/Celsr3^S1894Rfs*2/+^*(n=13/25), *p=0.0047*; Male *Celsr3^+/+^/Celsr3^S1894Rfs*2/+^*(n=11/16), *p=0.0042*. **(C)** Female *Wwc1^+/+^*/*Wwc1^W88C/+^*(n=17/20), *p=0.0066*; Male *Wwc1^+/+^*/*Wwc1^W88C/+^*(n=13/29) *p=0.3677*. **(D, E)** Aripiprazole attenuates PPI deficits in *Celsr3^C1906Y/+^* and *Celsr3^S1894Rfs*2/+^* females. **(D)** [F(11, 147)=6.491, *p<0.00001*, *R^2^*=*0.28*], genotype (*t*= −1.69, *p=0.093*), treatment (*t*=0.776, *p=0.44*), genotype (x) treatment (*t*=2.419, *p=0.017*). Vehicle-treated *Celsr3^+/+^*/*Celsr3^C1906Y/+^*(n=16/11); drug-treated *Celsr3^+/+^*/*Celsr3^C1906Y/+^* (n=13/13). € [F(11, 168)=6.987, *p<0.00001*, *R^2^*=*0.27*], genotype (*t*= −1.868, *p=0.064*), treatment (*t*=0.463, *p=0.64*), genotype (x) treatment (*t*=2.507, *p=0.013*). Vehicle-treated *Celsr3^+/+^*/*Celsr3^S1894Rfs*2/+^*(n=18/16); drug-treated *Celsr3^+/+^*/*Celsr3^S1894Rfs*2/+^*(n=13/13). (A, B, C) Two-way ANOVA, main effect genotype. (D, E) Multiple linear regression (factor III). *\*\*p<0.01, ***p<0.001*, *\*\*\*\*p<0.0001*. Full description of statistics in table S1.

### PPI deficits in Celsr3 female mice are attenuated by aripiprazole

We tested if PPI deficits in female *Celsr3^C1906Y/+^* and *Celsr3^S1894Rfs*2/+^* mice were responsive to aripiprazole, a partial agonist at dopamine type II and serotonin 5-HT1a receptors that is prescribed for TD. For the former group, we did not see a significant effect of treatment on levels of inhibition (*p=0.44*). We observed a trend-level effect of genotype (*p=0.093*) and a significant treatment (x) genotype interaction (*p=0.017*) For the latter, we also did not see a significant effect of treatment on levels of inhibition (*p=0.66*). We once again observed a trend-level effect of genotype (*p=0.064*) and a significant treatment (x) genotype interaction (*p=0.013*) (**Fig. 2D, E, Table S1**). Thus, PPI deficits appear responsive to aripiprazole, suggesting perturbations to dopamine and/or serotonin may contribute to sensorimotor gating deficits in *Celsr3^C1906Y^*^/+^ and *Celsr3^S1894Rfs^*^*2/+^ mice.

### *Celsr3* mutants perform better on an accelerating rotarod

Mice were tested on an accelerating rotarod to assess changes to motor coordination and balance (**Fig 3A, Table S1)**. *Celsr3^C1906Y/+^* mice displayed a longer latency to fall compared with wild-type littermates [males F(1, 30)=6.673, *p=0.015)*; females F(1, 32)=3.921, *p=0.056*]. *Celsr3^S1894Rfs*2/+^* males also showed a longer latency to fall [F(1, 41)=4.022, *p=0.051*], but *Celsr3^S1894Rfs*2/+^*females showed no differences compared to littermate controls. We did not detect changes to rotarod performance for *Wwc1^W88C/+^* mice (**Fig. 3A**). These results show that motor coordination and balance is improved in *Celsr3* mutants, and may also suggest changes to motor learning and execution.

**Figure 3:**
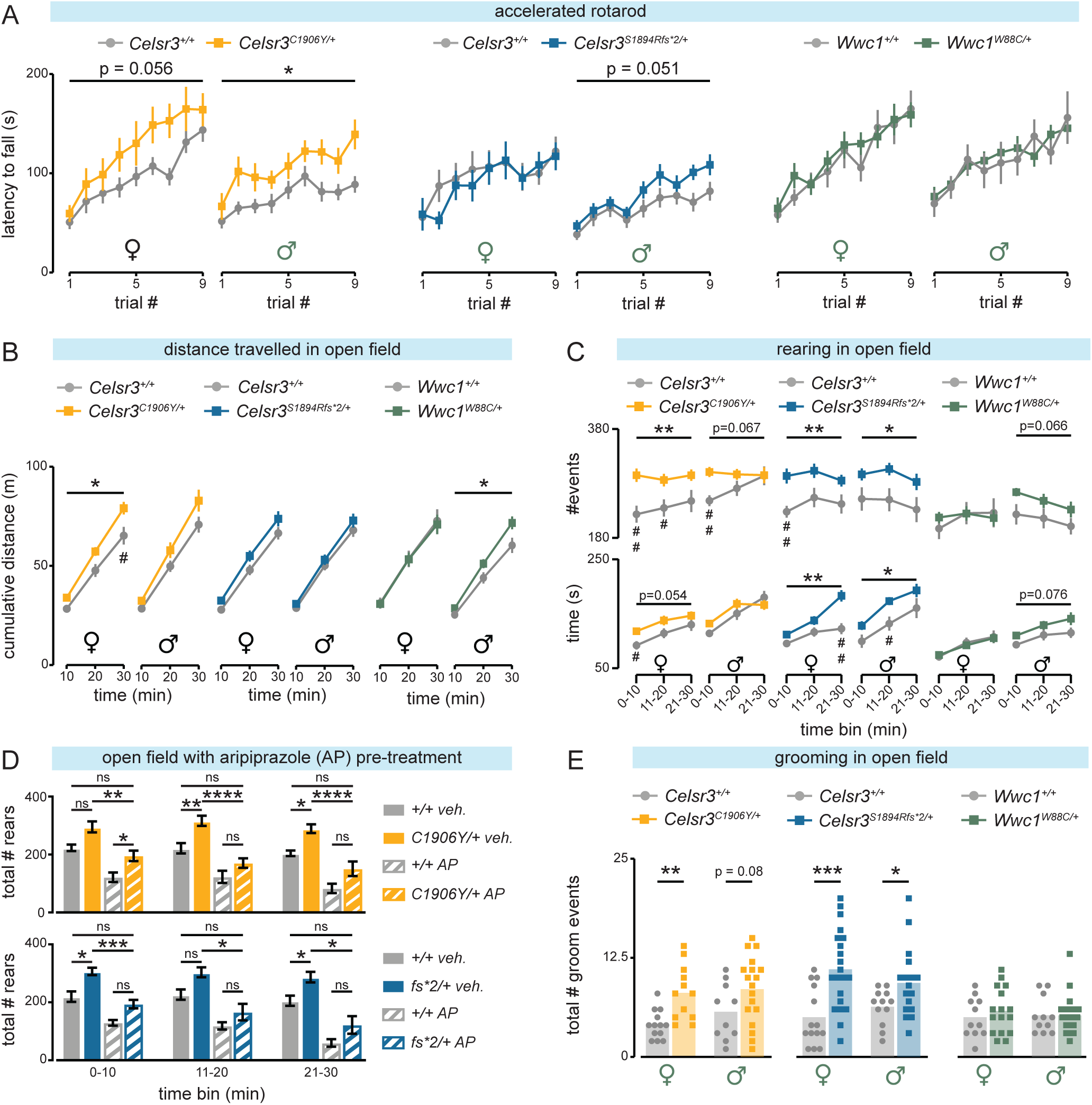
Mutant mice display improved rotarod performance with enhanced rearing and grooming in the open field. **(A)** *Celsr3^C1906Y/+^*males/females and *Celsr3^S1894Rfs*2/+^* males perform better on an accelerating rotarod. *Wwc1^W88C/+^* mice show normal latency to fall. **(B)** *Celsr3^C1906Y/+^*females travel more distance in the open field. *Celsr3^C1906Y/+^* males show a trend-level effect (*p=0.104*). *Celsr3^S1894Rfs*2/+^* females show a trend-level effect for more distance traveled (*p=0.102*) but males show no difference. *Wwc1^W88C/+^*males travel more distance in the open field but females travel similar distance. **(C**, Top Panel) Celsr3 males and females rear more in the open field. Rearing events trended up for *Wwc1^W88C/+^* males (*p=0.066*) but not females. (Bottom Panel) Time spent rearing within separate binned 10-min intervals. **(D)** Aripiprazole pretreatment attenuates rearing behavior. Wild-type/*Celsr3^C1906Y/+^*: [F(11, 114)=17.99, *p<0.00001*, *R^2^*=*0.59*], genotype (*p<0.0001*), treatment (*p<0.0001*), genotype (x) treatment interaction (*p=0.291*); Wild-type/*Celsr3^S1894Rfs*2/+^*: [F(11, 117)=18.13, *p<0.00001*, *R^2^*=*0.59*], genotype (*p<0.0001*), treatment (*p<0.0001*), genotype (x) treatment interaction (*p=0.269*) **(E)** *Celsr3^C1906Y/+^*females groom more in the open field and males show a trend. *Celsr3^S1894Rfs*2/+^* females and males also groom more. *Wwc1^W88C/+^* mice show no differences. **(A, B, C)** RM Two-way RM ANOVA, main effect genotype with Bonferonni’s Multiple Comparisons Test. **(A)** *Celsr3^+/+^*/*Celsr3^C1906Y/+^* female (n=20/14), male n**=**17/15; *Celsr3^+/+^*/*Celsr3^S1894Rfs*2/+^* female (n=20/18), male (n**=**17/29); *Wwc1^+/+^*/*Wwc1^W88C/+^*female (n=16/31), male (n=19/40). **(B, C)** *Celsr3^+/+^*/*Celsr3^C1906Y/+^* female (n=16/16), male n**=**15/17; *Celsr3^+/+^*/*Celsr3^S1894Rfs*2/+^* female (n=18/25), male (n**=**12/29); *Wwc1^+/+^*/*Wwc1^W88C/+^*female (n=12/21), male (n=13/18). **(D)** Multiple linear regression (factor III) with Tukey’s post-hoc test. Vehicle-WT/*Celsr3^C1906Y/+^*(n=12/10), Drug-WT/*Celsr3^C1906Y/+^* (n=10/10); Vehicle-WT/*Celsr3^S1894Rfs*2/+^*(n=12/11), Drug-WT/*Celsr3^S1894Rfs*2/+^* (n=10/10). **(E)** Mann-Whitney Test. *Celsr3^+/+^*/*Celsr3^C1906Y/+^* female (n=13/12), male (n**=**10/18); *Celsr3^+/+^*/*Celsr3^S1894Rfs*2/+^*female (n**=**14/21), male (n**=**12/17); *Wwc1^+/+^*/*Wwc1^W88C/+^* female (n=11/16), male (n=11/29) \**p<0.05*, *\*\*p<0.01, ***p<0.001*, *\*\*\*\*p<0.0001*; ^#^*p<0.05*, ^##^*p<0.01*. Full description of statistics in table S1.

### *Celsr3* mutants are more active and show enhanced rearing behavior in the open field

We assessed behavior while mice navigated an open field arena for 30 minutes (**Table S1**). *Celsr3^C1906Y/+^* females were more active and traveled more total distance compared to controls [F(1, 30)=6.110 *p=0.019*], whereas *Celsr3^C1906Y/+^* males showed a trend-level effect [F(1, 30)=2.814 *p=0.10*] (**Fig. 3B**). Latency to enter the center of the arena was similar for *Celsr3^C1906Y/+^* mice compared to controls (**Fig. S3A)**. *Celsr3^C1906Y/+^* females made more center zone entries over time [Time x genotype effect: F(2, 60)=5.693, *p=0.005*], whereas males showed a weak trend [Time x genotype effect: F(2, 60)=2.04, *p=0.139*] (**Fig. S3B**). Thus, *Celsr3^C1906Y/+^* mice, particularly females, are more active in the open field. Exploratory behavior may also be increased in females, as reflected by the greater number of center zone entries as time progressed.

Next, we looked at the number of vertical rearing events and time spent rearing. The cumulative number of rearing events was increased for *Celsr3^C1906Y/+^* females [F(1, 30)=9.1, *p=0.003*], as was total time spent rearing [F(1, 30)=4.374, *p=0.045*] (**Fig. S4A, B**). Rearing events also showed significant or near-significant increases when analyzed separately within binned intervals (0-10 min, *p=0.002*; 10-20 min, *p=0.046*; 20-30 min, *p=0.074*) (**Fig. 3C**). However, except for the first 10 minutes, time spent rearing within these intervals was not significantly different (0-10min, *p=0.045*; 10-20 min, *p=0.22*; 20-30 min, *p=0.73)* (**Fig. 3C**). This suggests some vertical rears may have been briefer and more repetitive-like. *Celsr3^C1906Y/+^*males displayed a trend for more rearing events across binned intervals [F(1, 30)=3.6, *p=0.067*)] (**Fig. 3C**). Rearing events were especially increased during the first 10 minutes (0-10min, *p=0.005*), however time spent rearing was not different (*p=0.2*) **(Figs. 3C and S4B).** This suggests *Celsr3^C1906Y/+^* males may also show some repetitive-like rearing. For males and females, rearing events along the perimeter of the arena were increased (females, *p=0.001* and males, *p=0.023*) (**Fig. S4C**).

*Celsr3^S1894Rfs*2/+^* females also traveled more distance, showing a trend-level effect, whereas males did not [Female: F(1, 41)=2.803, *p=0.1,* Male: F(1, 39)=0.689, *p=0.439*] (**Fig. 3B**). Latency to enter the center of the arena was shorter for *Celsr3^S1894Rfs*2/+^* females (*p=0.073*), whereas males showed no differences versus controls **(Fig. S3C).** *Celsr3^S1894Rfs*2/+^* females also showed a trend-level effect for more center zone entries [F(1,41)=3.61, *p=0.065*], but no changes were observed for *Celsr3^S1894Rfs*2/+^* males [F(1,39)=0.1922, *p=0.664*] **(Fig. S3D)**. The cumulative number of rearing events [F(1, 41)=10.8, *p=0.002*] and total time spent rearing [F(1, 41)=8.09 *p=0.007*] were also greater for *Celsr3^S1894Rfs*2/+^*females (**Fig. S4D, E**). Significant or near-significant differences in the number of rearing events were seen within binned 10-minute intervals (0-10 min *p=0.004*, 11-20 min *p=0.043*, and 21-30 min *p=0.098*) (**Fig. 3C)**. However, time spent rearing was similar to controls within the first and second intervals, similar to *Celsr3^C1906Y/+^* females (0-10 min *p=0.13*, 10-20 min *p=0.22*) **(Fig. 3C)**. The cumulative number of rearing events [F(1,39)=4.618, *p=0.038*] and total time spent rearing [F(1,39)=4.998, *p=0.031*] were also increased for *Celsr3^S1894Rfs*2/+^* males, but to a lesser extent versus females (**Fig. S4D, E)**. The number of rearing events was also significantly increased when measured separately within and across binned intervals [F(1, 39)=5.8, *p=0.021*] (**Fig. 3C**). Rearing events along the perimeter and center both trended upward for *Celsr3^S1894Rfs*2/+^* mice (**Fig S4F).** Thus, like *Celsr3^C1906Y/+^* mutants, *Celsr3^S1894Rfs*2/+^*males–and especially females–rear more in the open field.

*Wwc1^W88C/+^* males traveled more distance in the open field [F(1,29)=4.27, *p=0.049*], with differences becoming more apparent over time [time (x) genotype, F(2, 58)=5.5, *p=0.006*]. *Wwc1^W88C/+^* females, by contrast, traveled similar distances versus controls **(Fig. 3B)**. Latency to enter the center of the arena and number of entries was unaffected for both sexes **(Fig. S3E, F).** *Wwc1^W88C/+^*males also showed more cumulative rearing events [F(1,29)=4.2, *p=0.048*] and a trend for increased time spent rearing [F(1, 29)=3.6, *p=0.06*] **(Fig. S4G, H**). When measured across binned intervals, we saw trend-level effects for the number of rearing events [F(1,29)=3.6, *p=0.066*] and time spent rearing [F(1,29)=3.4, *p=0.076*] **(Figs. 3C)**. We did not observe preference for rearing along the perimeter versus the center (**Fig. S4I)**. Rearing behavior was normal for *Wwc1^W88C/+^* females, in stark contrast to *Celsr3* female mutants (**Figs. 3C, S4G-I**). Thus, *Wwc1^W88C/+^* males are slightly more active and rear more in the open field, while females behave similarly to controls.

### Aripiprazole attenuates rearing behavior in the open field

We tested if aripiprazole pretreatment could attenuate rearing behavior for *Celsr3^C1906Y/+^* and *Celsr3^S1894Rfs*2/+^*mice, focusing on females who showed more pronounced changes. We observed significant effects of genotype (*p<0.0001*) and treatment (*p<0.0001*) on rearing events for *Celsr3^C1906Y/+^* and also *Celsr3^S1894Rfs*2/+^* mice (genotype, *p<0.0001*; treatment, *p<0.0001*) (**Fig. 3D**). Post-hoc comparison testing showed rearing counts were significantly increased for vehicle-treated mutants (both genotypes) versus vehicle-treated controls (**Table S1**). Aripiprazole significantly reduced the number of rearing events for drug-treated versus vehicle-treated controls, as well as for drug-treated versus vehicle-treated mutants (both genotypes). However, differences were no longer significant for vehicle-treated controls versus drug-treated mutants (both genotypes) (**Fig. 3D**). Thus, enhanced rearing behavior in mutant *Celsr3* females is attenuated by aripiprazole.

### Grooming events are increased for *Celsr3* mutants

The number of grooming bouts, total time spent grooming, and average bout duration were measured in the open field (**Table S1**). The total number of grooming bouts was increased for *Celsr3^C1906Y/+^ (p=0.001)*) and *Celsr3^S1894Rfs*2^* females (*p=0.0005*) (**Fig. 3E**). Hair loss was sometimes found behind the ears and back of the neck, but no lesions **(Fig. S5A)**. Total time spent grooming was also increased for females (*Celsr3^C1906Y/+^*, *p=0.052*; *Celsr3^S1894Rfs*2/+^ p=0.001*)) **(Fig. S5B, C)**. The average duration of each grooming bout was comparable to controls, suggesting sequential, stereotypic grooming sequences are not interrupted (**Fig. S5B, C**). The number of grooming bouts was also increased for both *Celsr3^C1906Y/+^* and *Celsr3^S1894Rfs*2/+^*males (*p=0.082* and *p=0.045*, respectively) (**Fig. 3E)**. We did not detect significant changes to total time spent grooming nor the average duration of each grooming event, although the latter trended shorter in *Celsr3^C1906Y/+^* and *Celsr3^S1894Rfs*2/+^* males **(Fig. S5B, C)**. The number of grooming bouts, total time spent grooming, and average grooming bout duration were normal in male and female *Wwc1^W88C/+^* mice **(Figs. 3E, S5D)**. Repetitive digging behaviors were also assessed using the marble burying assay. We did not observe any significant enhancement for either *Celsr3* and *Wwc1* mutants **(Fig. S6)**. Thus, increased grooming, in addition to rearing, constitute motor phenotypes that are prominent in *Celsr3^C1906Y/+^* and *Celsr3^S1894Rfs*2/+^* females but, interestingly, less so in males.

### Unsupervised machine learning reveals changes to spontaneous motor behavior

We characterized changes to spontaneous motor behavior using Motion Sequencing (MoSeq). MoSeq uses 3D depth imaging and unsupervised machine learning to parse motor behavior into the sequential expression of reusable and stereotyped behaviors (i.e., syllables) while mice navigate a circular open field (25) (**Fig. 4A**). These syllables typically last a few hundred milliseconds and estimates for syllable usage frequency and duration can be measured. Further, MoSeq can assess how behavior is organized through analysis of transition probabilities and syllable sequence entropy (i.e., how mice arrange syllables into patterned movement sequences across time), advantageous for movement disorders like TD. Co-training MoSeq with data from each model revealed numerous changes to syllable usage frequencies in mutant mice. In male and female *Celsr3^C1906Y/+^*and *Celsr3^S1894Rfs*2/+^* mice, shared syllables corresponding to rearing (both assisted and unassisted) were generally upregulated (**Fig. 4Bi-vi)**). Conversely, syllables corresponding to pauses were generally downregulated, perhaps reflecting their overall tendency to be more active. Changes to syllable usages were less prominent in *Wwc1^W88C/+^* mice, and did not substantially overlap with Celsr3 mutant mice (**Figs. 4v-vi, Table S2**).

**Figure 4:**
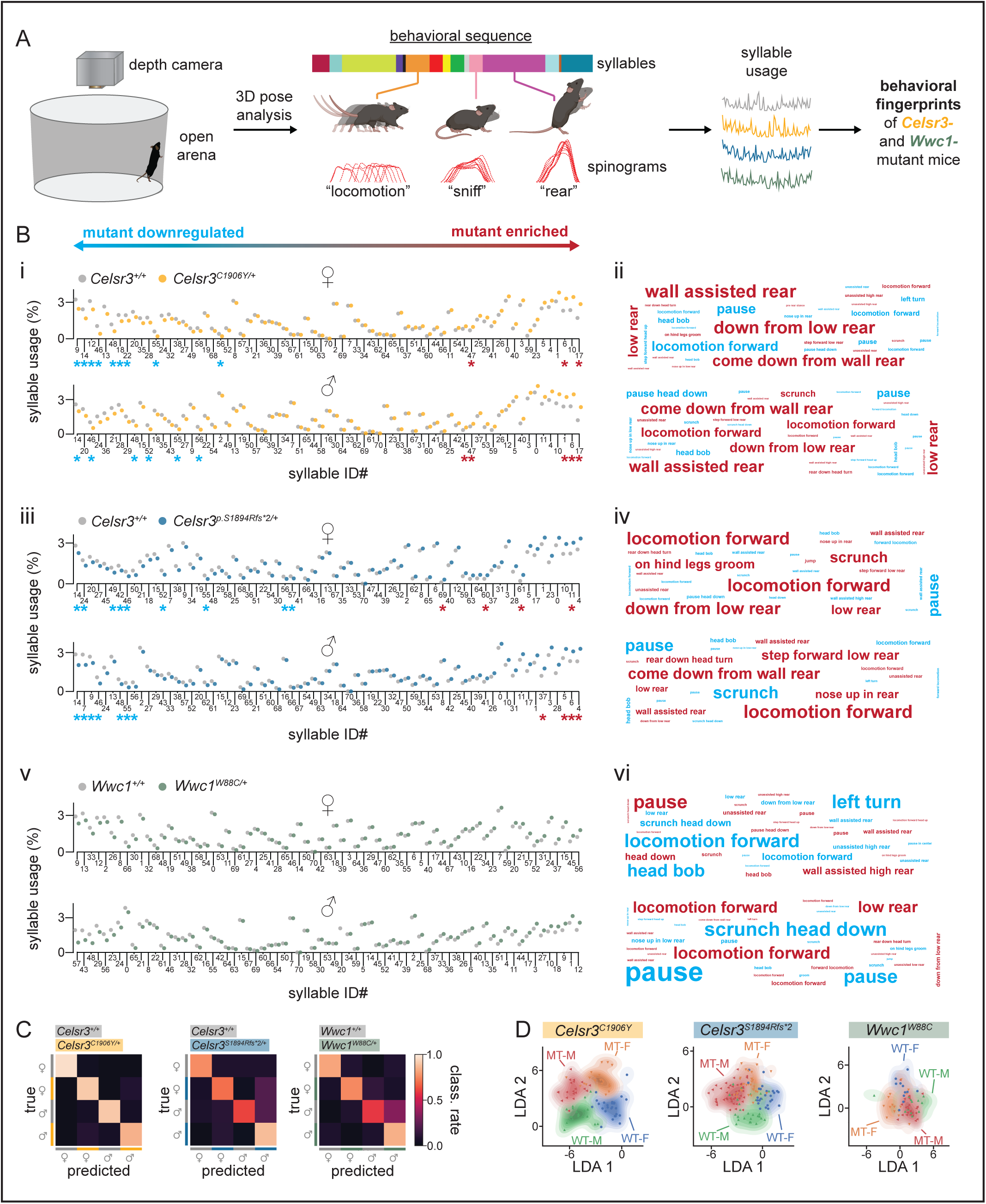
Unsupervised machine learning reveals changes to spontaneous motor behavior. **(A)** Schematic of motion sequencing approach. **(B)** Left, mutation plots summarizing mean usage of each syllable in wild type and mutant mice. Syllables are ordered by the relative difference between mutants and wild-type (left: mutant downregulated; right: mutant enriched). Syllables with significant changes in usage, identified by Mann-Whitney U test and post hoc Benjamini-Hochberg correction, are indicated by asterisks (*p<0.05*). Right, word clouds representing relative syllable changes in mutants compared to controls. Word color indicates direction of change (red: mutant upregulated; blue: mutant downregulated). Word size is proportional to the difference in usage between mutants and controls. Words are ethological descriptors assigned by reviewers. **(C)** Normalized classification matrices showing the performance of a classifier trained on syllable usage for mice grouped by line, sex, and genotype. An ideal classification is a value close to 1, shown in white. **(D)** Linear discriminant analysis plots showing similarity of mutant and wild type groups based on syllable usage signatures. Full description of statistics in table S2.

We used linear discriminant analysis (LDA) to learn a low-dimensional (2D) mouse embedding, trained on syllable usages, to better visualize and categorize mice across sex and genotype. The LDA projection for *Celsr3^C1906Y/+^*and *Celsr3^S1894Rfs*2/+^* males and females showed good separation by sex and genotype from their respective controls, whereas *Wwc1^W88C/+^*mice were less segregated from littermate controls **(Fig. 4C, D)**. We found that these models could predict the sex and genotype of held-out mice with high accuracy, which was ablated by randomization of group labels, indicating that a true relationship exists between the genotype, sex of mice, and the syllable usage values they emit (*Celsr3^C1906Y^* model: accuracy on held-out data=0.56, permutation test *p=9.99e-4*; *Celsr3*^S1894Rfs*2^ model: accuracy on held-out data=0.85, permutation test *p=9.99e-4*) (**Fig. S7**). We also computed the cosine distance within- and between-groups to further probe group separability within the high-dimensional syllable usage space. The between-group cosine distance for all sexes and genotypes exceeded the within-group cosine distances in all cases **(Fig. S8**). This indicates that, overall, members of each group defined by mouse line, sex, and genotype are closer to each other than mice from another group in the high-dimensional syllable usage space.

Syllable usage entropy revealed mutant mice displayed lower entropy relative to controls (**Fig. 5A, left**). This indicates that spontaneous motor behavior is less diverse on average, which might be expected with increased expression of repetitive behaviors (e.g., rearing). However, syllable *sequence* entropy rate was increased, suggesting the transitions between motor sequences, reflecting motor planning or action selection strategies that regulate patterned movements across time, were more disorganized and/or unpredictable (26) (**Fig. 5A, right**). When comparing state maps of relative changes to transition probabilities between mutants and controls, some common altered transitions between certain syllables are apparent across multiple mutant lines (**Figs. 5B, S9A-C**). Altogether, these findings suggest *Celsr3* mutant mice favor certain behaviors at expense of others, but the action selection strategies to reach these more favored states become less predictable. Additionally, mutant mice are discriminated from normal behaving controls according to their respective syllable usage frequencies and transition probabilities.

**Figure 5:**
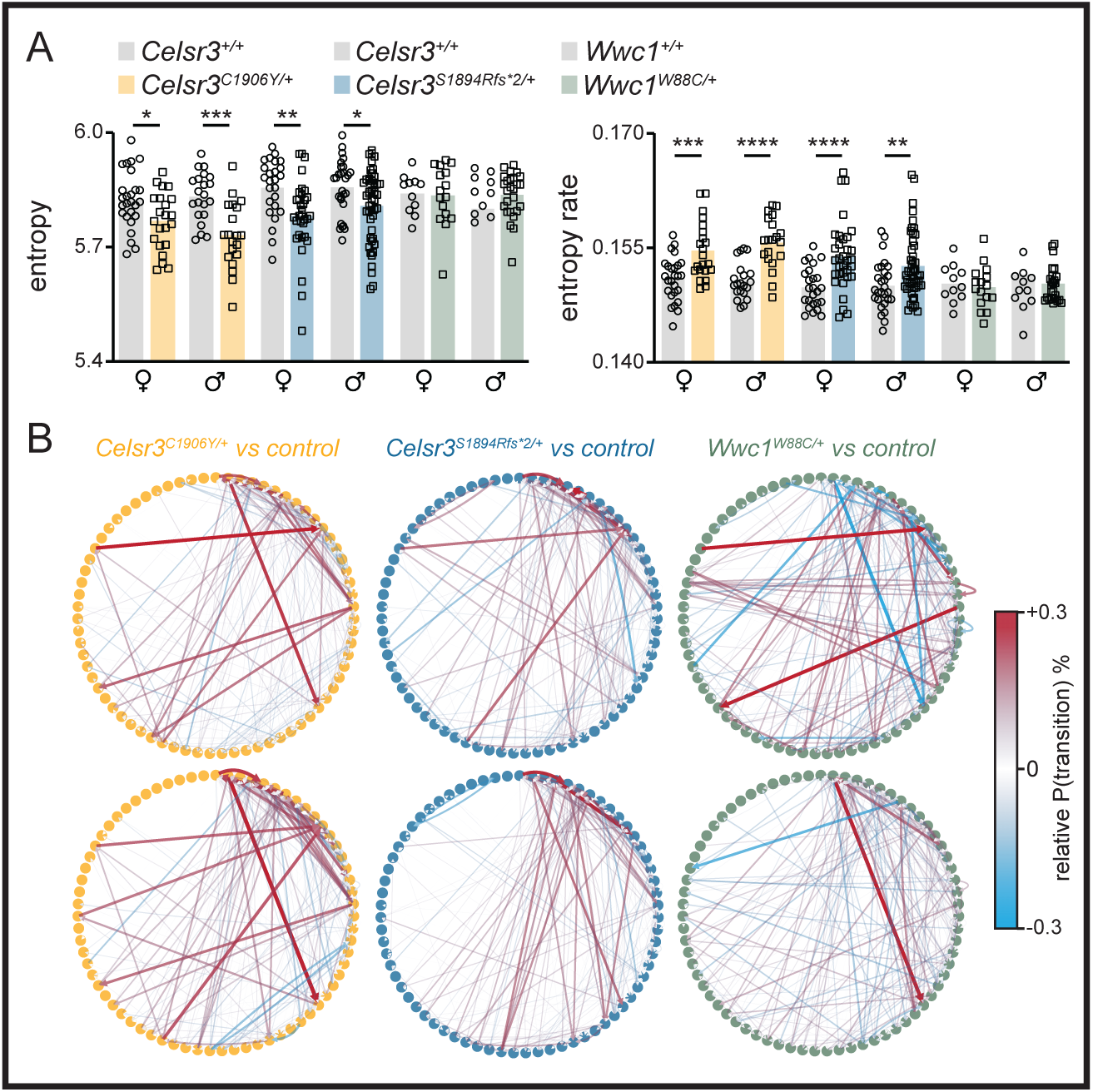
Motion sequencing shows common changes to behavioral structures in mutant mice. **(A)** Left, entropy graph. Right, entropy rate graph. **(B)** State maps showing transition probability changes in mutants relative to wild type mice. Circles represent syllables, lines represent transitions with red lines showing a higher relative transition probability and blue lines showing a lower relative transition probability. *Celsr3^C1906Y/+^*females (top left, *Celsr3^+/+^* n=26, *Celsr3^C1906Y/+^* n=22) and males (bottom left, *Celsr3^+/+^* n=21, *Celsr3^C1906Y/+^* n=20). *Celsr3^S1894Rfs*2/+^*, females (top middle, *Celsr3^+/+^* n=25, *Celsr3^S1894Rfs*2/+^*n=33) and males (bottom middle, *Celsr3^+/+^* n=28, *Celsr3^S1894Rfs*2/+^*n=48). *Wwc1^W88C/+^* females (top right, *Wwc1^+/+^*n=11, *Wwc1^W88C/+^* n=16), and males (bottom right, *Wwc1^+/+^*n=12, *Wwc1^W88C/+^* n=26).

### Reward learning and evoked dopamine release is enhanced in *Celsr3* mutants

Reward learning is suggested to be altered in humans with TD but, to our knowledge, has not been investigated in animal models (27). Given changes to sensorimotor behavior seen in *Celsr3* mutant mice, we next wanted to assess whether goal-driven behavior was also affected. We examined motor responding and reward learning in *Celsr3^C1906Y/+^* males and *Celsr3^S1894Rfs*2/+^*females (consistent with the sex of the affected human proband) using a continuous, fixed-ratio reinforcement schedule (**Table S1**). In this assay, repeated over a span of four days, mice have ninety minutes to earn a total of thirty sucrose pellets, with one pellet earned for each nose poke. Upon examining rewarded nose poke timestamps across individual trials, *Celsr3^C1906Y/+^* and *Celsr3^S1894Rfs*2/+^*mice showed elevated motor responding compared to wildtype littermates (**Fig. 6A**). The rate of cumulative rewards obtained was significantly enhanced on all four days of testing for *Celsr3^C1906Y/+^* and *Celsr3^S1894Rfs*2/+^* mice (*p<0.0001*), as determined by linear (day 1, R^2^ > 0.99) and non-linear regression analysis (days 2-4, R^2^>0.98) (**Fig. 6B**). The mean time spent to obtain thirty rewards was reduced in *Celsr3^C1906Y/+^* mice on day three (*p=0.03*) and in *Celsr3^S1894Rfs*2/+^* mice on days two (*p=0.04*) and three (*p=0.03*) (**Fig. 6C, top**). Analyzed across all four days, mean time to completion was significantly shorter [*Celsr3^C1906Y/+^*, F(1, 219)=4.998, *p=0.026*; *Celsr3^S1894Rfs*2/+^*, F(1, 191)=11.43, *p=0.0009*]. Additionally, across days two through four, we saw a trend-level effect for reduced latency to retrieve first rewards, suggesting motivation might be increased **(Fig. 6C, bottom)**. We next measured changes to electrically evoked striatal dopamine using fast scan cyclic voltammetry. *Celsr3^C1906Y/+^* mice have higher peak evoked DA in the dorsal striatum compared to controls (*p=0.002*) (**Fig. 6D-I**). The decay constant was normal (**Fig. 6J**). Thus, motor responding and reward learning are enhanced in *Celsr3* mutants, potentially due to effects on striatal dopamine.

**Figure 6:**
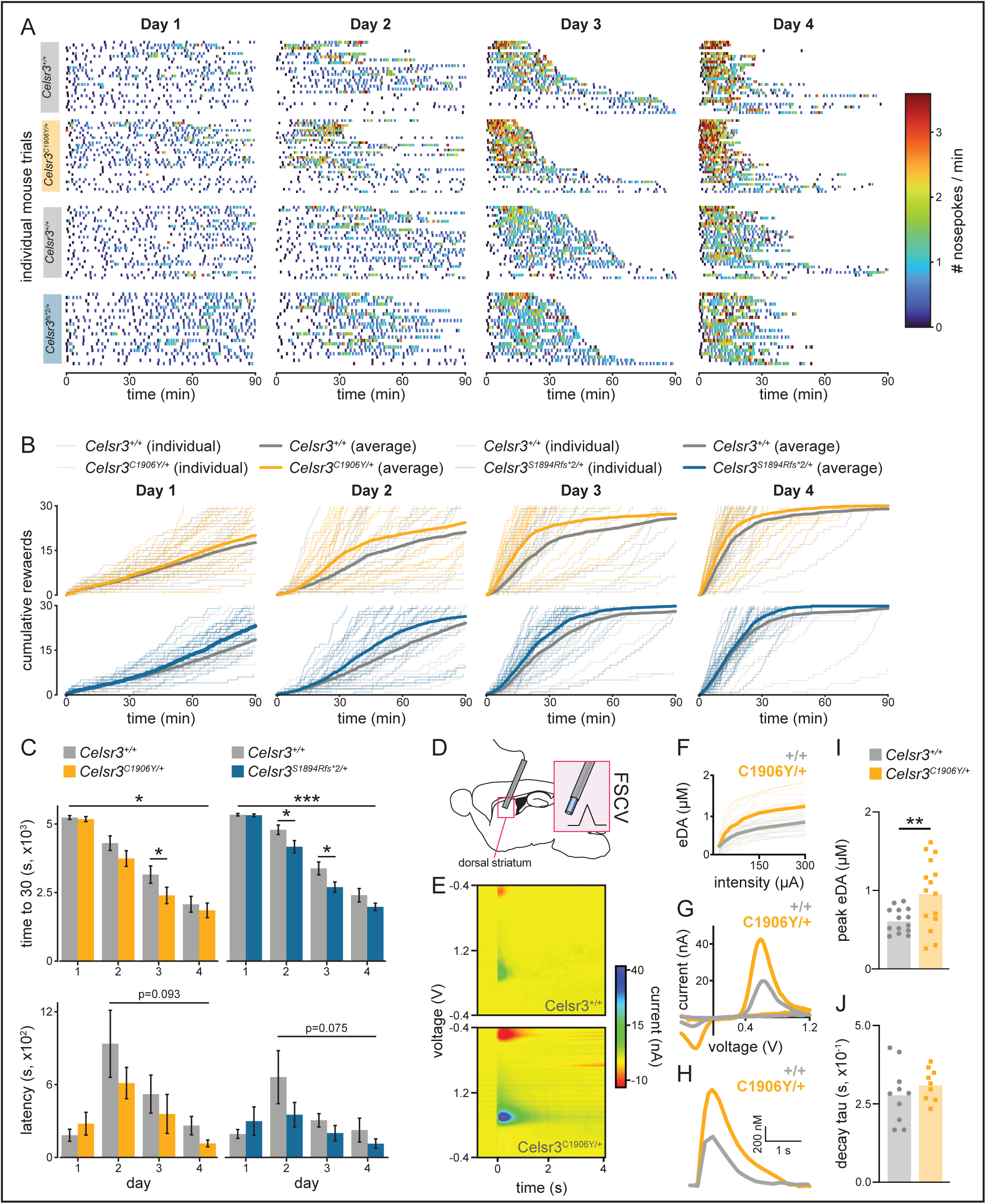
*Celsr3^C1906Y/+^ and Celsr3^S1894Rfs*2/+^* mice show enhanced motor responding and reward learning. Male *Celsr3^+/+^* (n=26), *Celsr3^C1906Y/+^*(n=30), and female *Celsr3^+/+^* (n=27), *Celsr3^S1894Rfs*2/+^*(n=22) mice were trained in a fixed ratio-1 (FR1) paradigm (one nose poke=one pellet). **(A)** Individual trials with nose poking events color-coded to indicate instantaneous rate. **(B)** Cumulative number of rewards earned during 90-minute sessions across training days. Thin transparent lines represent individual mouse performances. Bold lines show mean cumulative rewards across one second bins for mutants (orange=*Celsr3^C1906Y/+^*, blue=*Celsr3^S1894Rfs*2/+^*) or controls (gray=*Celsr3^+/+^*). The rate of cumulative rewards earned is significantly different on all days in mutants vs. wildtypes [*p<0.05*, linear regression (day 1), non-linear regression (days 2-4)]. **(C)** Time required to reach 30 rewards (defaulting to 90 minutes if failed) graphed over four days. *Celsr3^C1906Y/+^* mice have a faster completion time on day 3 (*p=0.027*), and *Celsr3^S1894Rfs/+^* mice have a faster completion time on days 2 (*p=0.041*) and 3 (*p=0.029*, Mann-Whitney U tests). Across all four days, time is significantly shorter for mutant mice (*Celsr3^C1906Y/+^ p=0.026*; *Celsr3^S1894Rfs*2/+^*, *p=0.0009*). A trend-level effect for reduced latency to retrieve first reward is observed across days two-four (*Celsr3^C1906Y/+^*, *p=0.093*; *Celsr3^S1894Rfs*2/+^*, *p=0.075*). **(D)** Fast-scan cyclic voltammetry was performed to measure dopamine release in the dorsal striatum. **(E)** Representative 3D voltammograms showing background-subtracted current (false color scale) as a function of time (x) and voltage applied (y). **(F)** Evoked DA (eDA) relative to stimulation intensity, individual trials shown as fine lines and group averages shown as bold lines [genotype x stimulation intensity F(7,182)=4.489, *p=0.0001*, main effect genotype F(1,26)=7.393, *p=0.01*]. **(G)** Characteristic current peaks at 0.6 V and −0.2 V match the oxidation and reduction potentials for DA. **(H)** Representative eDA waveforms of *Celsr3^+/+^* (n=3) and *Celsr3^C1906Y/+^* (n=4) mice. **(I)** *Celsr3^C1906Y^* mice have higher peak eDA in the striatum (unpaired t test, *p=0.002*). **(J)** No differences were seen in the eDA decay time constants (tau). \**p<0.05*, *\*\*p<0.01, ***p<0.001*

### Brain development is grossly normal without signs of striatal interneuron loss

Celsr3 is required for axon pathfinding and the development of forebrain commissures (20). Cortical and striatal interneuron migration defects have also been reported in homozygous *Celsr3^GFP^* mice, in which GFP is knocked into the endogenous locus, ablating protein functions (14). We looked for signs of striatal interneuron loss in *Celsr3* and *Wwc1* mutant mice. We did not detect changes in density or nearest neighbor distance for parvalbumin-expressing interneurons (**Fig. 7A-F**). Crossing *Celsr3^S1894Rfs*2/+^*and *Celsr3^C1906Y/+^* mice with a transgenic reporter line that expresses green fluorescent protein (*eGFP*) in choline acetyltransferase (ChAT) expressing cells, or antibody labeling against ChAT in *Wwc1^W88C^* mice, did not reveal changes in density or nearest neighbor distance for striatal cholinergic interneurons (**Fig. 7A-F**). We also assessed overall cellular organization and development of major axon tracts in *Celsr3^S1894Rfs*2/+^*mice. Nissl staining showed grossly normal organization of cortical and subcortical regions (**Fig. S10A**). Crossing a genetic reporter line that expresses red fluorescent protein (*tdTomato*) in dopamine receptor type 1 (*Drd1a*) expressing striatonigral fibers did not reveal any major changes to axon fiber trajectory (**Fig. S10B**). Cortical layering, thickness, and within-layer cell density were normal, as determined by immunolabeling for transcription factors Ctip2, Satb2, and Foxp2 (**Fig. S10C**). Thus, brain development is grossly normal in *Celsr3^S1894Rfs*2/+^* mice, and striatal interneuron loss is not detected in all three models.

**Figure 7:**
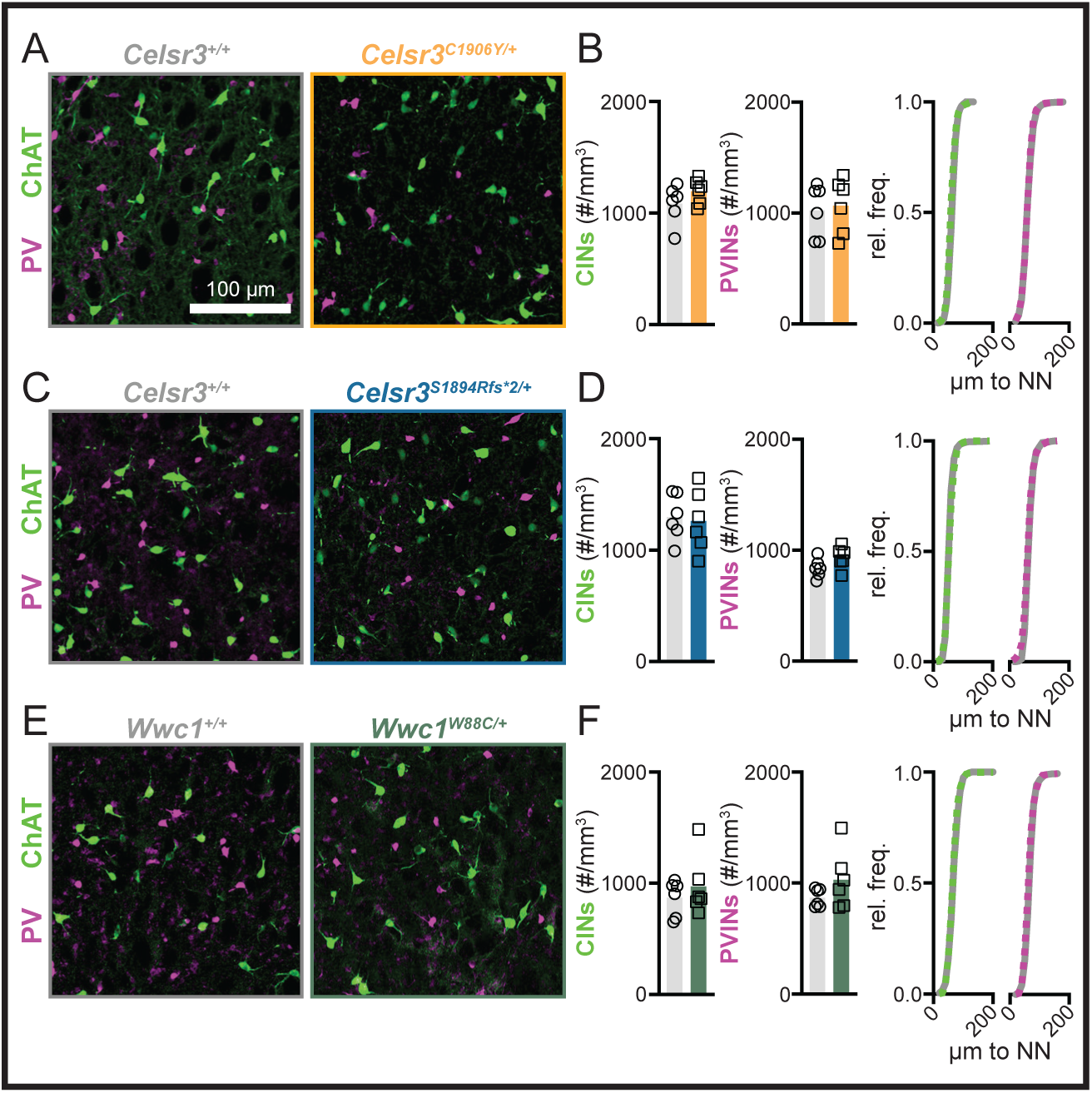
Striatal cholinergic and parvalbumin-expressing interneuron density is normal in *Celsr3* and *Wwc1* mutant mice. **(A, C, E)** Representative images showing striatal CINs (ChAT+, green) and PVINs (PV+, magenta) in *Celsr3^C1906Y/+^*, *Celsr3^S1894Rfs*2/+^*, and *Wwc1^W88C/+^* mice and littermate controls. **(B)** Quantification of CIN (left, *p*=0.2220) and PVIN density (middle, *p*=0.7735) in *Celsr3^+/+^* (n=6) and *Celsr3^C1906Y/+^* (n=6) mice, and cumulative frequency distribution of nearest neighbor (NN) distance of CINs and PVINs (right, *Celsr3^+/+^*: solid curve, *Celsr3^C1906Y/+^*: dashed curve). **(D)** Quantification of CIN (left, *p*=0.8090) and PVIN (middle, *p*=0.1000) density in *Celsr3^+/+^* (n=6) and *Celsr3^S1894Rfs*2/+^* (n=6) mice, and cumulative frequency distribution of NN distance of CINs and PVINs (right, *Celsr3^+/+^*: solid curve, *Celsr3^S1894Rfs*2/+^*: dashed curve). **(F)** Quantification of CIN (left, *p*=0.4579) and PVIN (middle, *p*=0.2077) density in *Wwc1^+/+^* (n=6) and *Wwc1^W88C/+^*(n=6) mice, and cumulative frequency distribution of NN distance of CINs and PVINs (right, *Wwc1^+/+^*: solid curve, *Wwc1^W88C/+^*: dashed curve). All statistical comparisons were made using Welch’s t tests.

## Discussion

Few genetic models have been developed for TD. Heterozygous *Ash1l* and homozygous histidine decarboxylase (*Hdc*) knockout mice possess validity at the gene level, although they do not model the exact human mutations (21, 28). Similar to our models, *Ash1l^+/-^* mice show elevated grooming whereas *Hdc^-/-^*mice have PPI deficits (28). However, PPI deficits have not been reported in *Ash1l^+/-^* mice and *Hdc^-/-^*mice do not show repetitive motor behaviors at baseline. Our testing suggests *Wwc1^W88C/+^* females may converge more closely with *Hdc^-/-^*mice, whereas *Celsr3^C1906Y/+^* and *Celsr3^S1894Rfs*2/+^*mice show behavioral phenotypes consistent with both *Ash1l^+/-^* and *Hdc^-/-^* mice, suggesting they provide distinct advantages for modeling TD. PPI deficits are the most consistent finding between ours and the aforementioned models, and may reflect a core behavioral phenotype in TD mice.

Despite detecting changes to motor behavior and cognitive functions, brain development was grossly normal in *Celsr3^S1894Rfs*2^*mice, and striatal interneuron loss was not detected in any of the three models. Furthermore, striatal interneuron loss is absent in more mildly affected *Celsr3^R774H/R774H^* mice reported elsewhere, in agreement with findings from *Hdc^-/-^* and *Ash1l^+/-^*mice (28–30). Notably, striatal interneuron loss was reported in postmortem tissue from a small cohort of adults with severe, refractory TD whose gene mutations, if any, were unknown (31, 32). A follow-up study using brain organoids grown from patient derived iPSCs (without identified gene mutations) also showed interneuron loss (33). These studies are widely cited to explain the neuropathology of TD and have motivated the development of animal models based on diphtheria-toxin mediated interneuron ablation or bicuculline disinhibition in the striatum (34–39). Although these studies have provided interesting insights, these putative models lack construct validity, do not recapitulate neurodevelopmental changes, and phenotypes can be mixed and/or subtle. Collectively, findings from multiple genetic models suggest striatal interneuron loss is rare in TD, perhaps limited only to severely affected adults.

Dopamine imbalances affecting fronto-cortico-striatal circuits are suspected in TD, despite limited direct evidence for DA involvement (40–48). Antipsychotic medications that act on dopamine D2 receptors (e.g., haloperidol and aripiprazole) are effective for reducing tics in some (41, 49). Dysregulated striatal dopamine is also suspected in *Hdc^-/-^* and *Ash1l^+/-^* knockout mice, with some repetitive motor behaviors responsive to haloperidol (21, 28). Certain behavioral phenotypes in *Celsr3^C1906Y/+^* and *Celsr3^S1894Rfs*2/+^*mice, including enhanced rearing and improved rotarod performance, are also suggestive of altered striatal dopamine (50, 51). Moreover, rearing behavior was attenuated by aripiprazole. Enhanced motor responding and reward learning under fixed reinforcement also suggest striatal dopamine levels may be higher. Indeed, electrically evoked dopamine release was increased in the dorsal striatum of *Celsr3^C1906Y/+^* mice. In addition to dorsal regions that control action-outcome learning and sensorimotor behavior, our results suggest affection to limbic regions, causing stronger motivational drive and/or reward perception (52–54). Higher levels of dopamine may enhance the perceived and/or anticipated rewarding effects of tics in response to premonitory urges. Over time, this may potentiate stimulus-response associations that produce the ‘urge-to-tic’ in certain environments, contexts, or emotional states. In this sense, tics may be maladaptive habits learned to alleviate unpleasant sensations and urges (55). Furthermore, dopamine stimulation and/or in vivo recordings paired with MoSeq show larger dopamine transients can reinforce syllable usages and ordering, promoting more variable behavioral sequences (i.e., higher sequence entropy rate) via effects on transition probabilities that govern syllable switching (56). Similarly, the observed changes we see with syllable usages, transition probabilities, and sequence entropy in Celsr3 mutants may be indicative of altered dopamine transients influencing the probability of performing certain syllables over others, and how these are ordered across space and time.

The prevalence of TD is higher in males than females (∼3-4:1), similar to autism spectrum disorder which shares some genetic overlap (6, 12, 57). While *Celsr3^C1906Y/+^* and *Celsr3^S1894Rfs*2/+^* males exhibit TD-like behaviors, in some instances the phenotypes appear stronger in females (e.g., hyperactivity, rearing, grooming), a somewhat unexpected finding. Moreover, *Wwc1^W88C/+^*females, but not males, have PPI deficits. Interestingly, a study in humans with OCD reported prepulse inhibition deficits only in females (58). It is still uncertain whether more common comorbidities in males, especially ADHD, may facilitate earlier diagnosis, skewing the true male/female sex ratio in TD (59). Some studies suggest sex bias in TD is less prominent in adults, and females may experience more compulsions, anxiety, and worsening of symptoms over time compared with males, who may be more likely to experience tic contraction (59–61). Our studies were limited to adult mice, and thus behavioral phenotypes could be overrepresented in adult females versus juveniles. Interestingly, damaging *de novo* variants are enriched in females from TD/CTD simplex families, especially in mutation intolerant genes such as *Celsr3* (12). This may explain why some phenotypes appear more penetrant in female mice. *Celsr3^S1894Rfs*2/+^* was also identified in a female with early onset tics noted by age three. Thus, our genetic models may provide important insights into naturally occurring sex differences in TD, and caution against selecting models or biasing studies towards males.

In summary, the most consistent findings in our genetic models are sensorimotor gating deficits and elevated rearing behavior in the open field. These phenotypes are responsive to aripiprazole in females (males were not tested). Enhanced reward learning is a novel phenotype in TD mouse models and may reflect higher phasic dopamine release. Changes to spontaneous motor behavior and first order transition probabilities between syllables discriminate patterned movement sequences between freely behaving TD mice versus wild-type controls. Notably, motor syllables are associated with distinct neuronal ensembles in the striatum, which may be reinforced by dopamine fluctuations (26, 56). While there is no clear consensus on what tics look like in mice, leveraging these unbiased computational approaches could thus prove useful for high throughput drug screening. Furthermore, pairing our models with genetic dopamine sensors (e.g., dLight) while they behave in various contexts (e.g., reward learning) may reveal new insights into neuronal substrates that underlie TD.

## Materials and Methods

### Animals

Strains and testing are approved by Rutgers IACUC PROTO201702623 (MAT) and IACUC PROTO999900331 (JAT). Mice were maintained on pure C57BL6/J backgrounds (JAX #000664). F1 founders were backcrossed for at least six generations. Mice were maintained on normal light-dark cycles and tested during the light cycle. Mice were group housed with littermates and separated by sex, and provided food and water *ad libitum* unless otherwise specified. Commercially available strains include *Drd1a-tdTomato* (RRID:IMSR_JAX:016204) and *Chat^BAC^-eGFP* (RRID:IMSR_JAX:007902). Complete details regarding mouse generation, DNA editing, and genotyping can be found in supplemental methods.

### Mouse behavior

Prepulse inhibition was run as previously described using mice aged 8-12 weeks (21). For rotarod, six-week-old mice completed an accelerated rotarod test (Harvard Apparatus) consisting of three trials per day for three consecutive days. Open field activity of eight-week-old mice was measured in a novel open field arena with activity monitor software (Med. Associates, Env-520, legacy). For fixed reinforcement, male *Celsr3^C1906Y/+^*and female *Celsr3^S1894Rfs*2/+^*mice were tested in sound attenuated operant chambers (Med Associates) under a fixed ratio-1 schedule lasting 90 minutes (or max 30 rewards) for four consecutive days. Nose poke time stamps and latency to reward retrieval were registered using Med-PC V (Med Associates) and sessions were recorded. Complete descriptions for behavioral procedures can be found in supplemental methods.

### Motion Sequencing

Analysis was performed using standard methods (26) and as previously described (62). Mice, median age of eight weeks, were placed individually in an open field arena and allowed to freely behave for 20 minutes while imaged from above with a Kinect v2 depth camera. Raw depth video was extracted, projected onto a common reference mouse to account for size differences, and projected onto a learned PCA embedding and modeled with autoregressive hidden Markov models (AR-HMM, a.k.a “MoSeq model”) on the first 10 PCs using a kappa found via search. Normalized syllable usage frequencies were tested using bootstrap tests with multiple testing correction by Benjamini-Hochberg procedure. Entropy and entropy rate were computed using standard formulas and tested using Mann Whitney U test. 5-fold cross-validated 2D LDA models were trained with normalized usage frequencies and following best practices. Complete descriptions of analysis procedures are found in supplemental methods.

### Mouse histology and striatal interneuron counts

Mice (both sexes, one month old) were transcardially perfused with 4% paraformaldehyde (PFA). Interneuron density was quantified in Imaris (Bitplane). Complete descriptions of all procedures, antibodies, and reagents can be found in supplemental methods.

### Statistical Analyses

Statistical analysis, with the exception of Motion Sequencing, was performed using Graphpad Prism (Graphpad Software Inc., La Jolla, CA). PPI was analyzed using two-way ANOVA across all three prepulses. Spearman’s correlation coefficient was performed to determine if correlations existed between startle amplitude and prepulse inhibition. Rotarod was analyzed across all nine trials using two-way ANOVA with repeated measures. Marble burying was analyzed using Mann Whitney test. Time spent grooming, time/grooming bout, and number of grooming events were analyzed using Mann-Whitney Test. Open field distance traveled and rearing events were analyzed using two-way ANOVA with repeated measures and Bonferroni multiple correction testing. Unless otherwise noted, two-way ANOVA results are presented as the main effect of genotype. For reinforcement learning, Mann-Whitney tests were used to compare time to completion, and t-tests with Welch’s correction and Holm-Šídák’s multiple comparison correction were used to compare rewards obtained. For MoSeq behavioral usage and transition matrices, bootstrap analysis was performed followed by Benjamini-Hochberg multiple comparisons testing. LDA analysis was completed using scikit-learn implementation. A complete description of all statistics and results for behavioral experiments are found in supplemental methods and Tables S1 and S2.

## Acknowledgements

The authors acknowledge the Rutgers Genome Editing Facility (Peter Romanienko and Ghassan Yehia). Jaclyn Eisdorfer for discussions pertaining to raster plots and statistics. Biorender.com for mouse schematics in Figs. 4 and S9. Funding sources: Tourette Association of America (MAT), New Jersey Center for Tourette Syndrome (MAT, GAH, JAT), Brain Research Foundation (#BRFSG-2022-11 MAT), Robert Wood Johnson Foundation (#74260, MAT), National Institutes of Mental Health R01MH115958 (JAT, GAH), NIH/NIMH U24 MH068457 (JAT).

