## Supplemental Material for "Human mutations in high-confidence Tourette disorder genes affect sensorimotor behavior, reward learning, and striatal dopamine in mice"

**Supplemental Materials:**

Mouse development and gene editing: C57BL6/J embryos were microinjected with a mixture containing Cas9 protein (IDT), an sgRNA (MilliporeSigma), and a ssODN (IDT) which contained homology arms for introducing each variant. Founders were screened by PCR and digested with restriction enzymes (introduced in PAM change). Variant insertions in founder mice were confirmed by PCR and/or next generation sequencing (Azenta Life Sciences). The following sgRNA and donor oligo sequences were used (sequence change in bold lower case, PAM change in bold upper case): **Celsr3-C1906Y**, sgRNA GGAGGAGGGCATCAGGGTCTGG (PAM is underlined) and donor oligo sequence CTATTTCTCAGGTCCCCAGTGACCCCATACCATGTTC TCATGACAGTCCTCACCTGAATA**t**AGCCAAC**G**AGACCCTGATGCCCCTCCTCCTTACTGCT GGGGGGCAGGCCTCCCACGTGGAGCTGTTTTAC; **Celsr3-pS1894Qfs*2** sgRNA CCCCTCC TCCTTACTGCTGGGGG (PAM is underlined) and the donor oligo sequence was GTGAGCTGCAGGGCCTGAAAGTAAAACAGCTCCACGTGGGAGGCCTGCCCCCCA**g**GCAG TAAGGAGGAGGGGCATCAGGGTCTGGTTGGCTGTATTCAGGTGAGGACTGTCATGAGAAC ATG (sequence and PAM change are both bold lower case); **WWC1-W88C**, sgRNA TCGAGGATCCAAGGGTGCAATGG (PAM is underlined) and donor oligo sequence GCTTCCTGAGCCACCACCAGGTAATCCTTCAGCATGTGCTCCTGTTCCCGCCG**g**CATTGCA CCCTTGGATCCTCGATCTGAGTGGTTTCTGAAAAAGATCCCAAAGCAATGTGTCAGCCATA C (sequence and PAM change are both bold lower case);

For *Celsr3*$^{C1906Y}$, amino acid p.Cys1906 was mutated to tyrosine by replacing UGU (Cys) with UAU (Tyr). Codon CUG for *p*.Leu1903 was switched to CUC (Leu/L) synonymously to introduce a BsaI site for genotyping. *Celsr3*$^{pS1894Qfs*2}$ was developed by adding a 1-base pair cytosine insertion (c.5679dupC). The insertion creates a BglI site for genotyping. Genotyping primers for *Celsr3*$^{C1906Y}$ and *Celsr3*$^{pS1894Qfs*2}$ are the same: Celsr3-J 5' GGTTTACCAGGTGCTTCTCCTTCG-3' and Celsr3-K 5'- CACTCCCATGCCAACATGTACTTG-3'. Amplified products were digested using Bsa1-HF (NEB, Cat#R3733L) and BglI (NEB, Cat#R0143L). The wild-type allele is 246 base pairs for both strains. Following BsaI digestion, two band sizes of 151 and 95 base pairs are generated for *Celsr3*$^{C1906Y}$. Following BglI digestion, two band sizes of 120 and 126 base pairs are generated for *Celsr3*$^{pS1894Qfs*2}$. For *Wwc1*$^{W88C}$, mice were genotyped using PCR primers WWC1A TAAAATGACGAGTCTCTGTACATCATG and WWC1B CAGCAATGGAAGGTACTCACAGC. The wild-type amplicon is 337 base pairs. Following NgoMIV digestion, the transgenic band produces fragment sizes of 148 and 189 base pairs.

Quantification of Celsr3 and Wwc1 protein levels: Whole brain lysates were prepared using synper (ThermoFisher, cat no: 87793). Protein quantity was measured using Pierce BCA assay kit (ThermoFisher cat. no. A55864). *Celsr3:* Lysates were run 3-8% Tris-Acetate PAGE gel. Protein was transferred from the gel onto a 0.45 um PVDF membrane overnight at 40V in 4C using NuPage transfer buffer plus 10% methanol (ThermoFisher cat. No. NP00061). The membrane was blocked for one hour in 5% non-fat milk. The membrane was next incubated at room temperature for 2 hours while rotating in Celsr3 antibody (1) and β-actin antibody (Invitrogen, PA1-183). Following antibody incubation, the membrane was washed 6x for 5 minutes in 0.1% PBS/Tween-20 followed by a 30-minute incubation at room temperature in Goat-anti-guinea pig-HRP (Invitrogen Cat no. A18769) and goat-anti-rabbit-HRP antibody (genscript). Following incubation, the membrane was washed 6x for five minutes in 0.1% PBS/Tween-20 and imaged (Kindle Biosciences KwikQuant). For the *Wwc1* western: Lysates were run on a 4-12% bis- tris PAGE gel at 180 volts for one hour in MOPS running buffer. Protein was then transferred on ice at 90V for one hour onto a 0.22um PVDF membrane with MOPS running buffer plus 20% Methanol on ice. The membrane was blocked for one hour in 5% nonfat milk. The membrane was then incubated in anti-Wwc1 (Cell Signaling Technology #8774) and anti-Beta-Actin antibodies, and incubated overnight at 4C while rotating. Following overnight incubation, the membrane was washed 6x for 5 minutes with 0.1% PBS/ Tween-20 at room temperature. Following the washes, the membrane was incubated in secondary goat anti-rabbit-HRP for 30 minutes, rocking at room temperature. The membrane was washed 6x for 5 minutes with 0.1% PBST at room temperature and imaged (Kindle Biosciences KwikQuant). Area under the curve was used to measure protein band density (ImageJ).

**Mouse Behavioral Procedures**

Prepulse Inhibition of the Acoustic Startle Reflex: Mice aged eight to twelve weeks were placed into the startle chamber (SR-Lab, San Diego Systems) and allowed to acclimate for five minutes before the start of the first trial. Background noise within the chamber was set to 65 dB. After the acclimation period, mice were subjected to five types of trial: 120 dB startle pulse alone, no pulse, and three prepulse trial types (6, 12 and 16 dB above background) followed 100ms later by a 120 dB startle stimulus. The intertrial interval averaged 15 seconds ranging from 8-23 seconds. The prepulse stimulus was 20ms in length and the startle pulse was 40ms in length. There were three blocks of trials. The first block consisted of six trials of startle pulse alone. The second block had

52 trials with a pseudorandom order of startle pulse alone, prepulse followed by startle pulse, and no pulse trials. The third block consisted of six trials of startle pulse alone. For the aripiprazole rescue, mice were injected intraperitoneally with 1 mg/kg of aripiprazole (Sigma, SML0935) or vehicle (0.9% saline/1% Tween-80/10% DMSO) 1-hour prior to undergoing the same trials as described above. Statistical analysis was performed using two-way ANOVA and GraphPad Prism, or multiple regression (type III) analysis.

Rotarod: Mice aged six weeks were tested. Prior to the first trial, mice were trained at a constant speed of 4 RPM for 30 seconds on the rotarod apparatus (Harvard Instruments). After 30 minutes, mice were placed on an accelerating rotarod (4-40 RPM). Mice underwent three trials per day for three consecutive days for a total of nine trials. Each trial ended when the mouse fell off or reached 300 seconds. The mean latency to fall was used for analysis. Statistical analysis was performed using GraphPad Prism and two-way ANOVA with repeated measures.

Open Field Arena: Eight-week-old mice were allowed to habituate to the room for at least thirty minutes. Locomotor activity was measured in a novel open field arena (Med. Associates Env-520, legacy). The plexiglass arena was 40 cm X 40 cm in dimension. A central zone was defined as 32.5 cm X 32.5 cm. Mouse activity was measured via infrared beam breaks in the X, Y and Z axes. At the start of the test, each mouse was placed in the same corner of the arena and allowed to freely explore for 30 minutes. Activity Monitor Software (Med Associates) was used to determine the distance traveled, rearing time, and rearing events. Aripiprazole was injected intraperitoneally 1 hour prior to entrance into the open field. Minimally sedating dosages specific for each line were used. *Celsr3*$^{C1906Y/+}$ mice and their wild-type littermates were dosed with 0.3 mg/kg or vehicle (0.9% saline/1% Tween-80/5% DMSO). *Celsr3*$^{S1894Rfs*2/+}$ mice and their wild-type littermates were dosed with 0.5 mg/kg or vehicle.

Grooming: Mice with a median age of eight weeks were scored. The mice were placed in the arena and allowed to acclimate for 10 minutes. Scoring was performed manually post hoc for the entirety of the subsequent 10 minutes from RGB videos. Observers were masked to genotype and sex. Total time spent grooming, number of grooming events, and time per grooming event were scored. Grooming was scored manually and independently by two masked observers.

Marble Burying Task: Mice aged between eight to ten weeks were placed in a standard mouse polycarbonate box containing 5 cm of beta chip bedding with glass marbles set on top of the bedding in a 4 X 5 lattice. After 30 minutes the number of marbles buried were counted. Any marble more than 2/3 buried was scored as buried. Marbles were counted blind to genotype by two separate observers. The observers' scores were averaged. Statistical analysis was performed using GraphPad Prism and the Mann Whitney Test. Masked analysis was performed.

Fixed Reinforcement: Randomly grouped mice were allowed access to regular chow and water ad libitum. Four grams of sugar pellets (Bio-Serv, Cat#F05301) per mouse were additionally provided in the home cage daily for one week prior to fixed reinforcement day 1 to acclimate mice to the reward. After acclimation, mice were food restricted for five days to attain 85%-90% normal body weight. Mice behaved in sound attenuated operant chambers. Nose poke holes and port lights were situated on opposite walls to the food magazine (Med Associates, Cat#ENV-115C). One day before testing, mice were placed in chambers for 1 hour to consume five 20-mg sugar pellets (Bios-Serv, Cat#F05301) in the center port and 10 sugar pellets in the food magazine to introduce the mice to the testing environment. Throughout the duration of fixed reinforcement testing, the central port cue light was illuminated. Each successful nose poke delivered a sugar pellet, and a new trial started after mice entered the food magazine to retrieve the reward. Each session comprised 30 rewards or 90 minutes, whichever came first. Testing lasted four consecutive days. Nose poke time stamps and latencies were registered using Med-PC V (Med Associates) and sessions were video recorded. Following daily testing, mice were fed 1.5-2.5 grams of regular chow to maintain body weight.

**Mouse histology and interneuron counts**: Striatal cholinergic (CIN) and parvalbumin-expressing (PVIN) interneuron densities were measured across the entire striatum in 100 μm coronal sections (Bregma +0.26 mm). CINs and PVINs were co-labelled in all animals. All sections were pre-incubated for one hour in 10% normal donkey serum and 0.5% Tween in PBS. In *Celsr3*$^{C1906Y/+}$;*Chat-eGFP*, *Celsr3*$^{S1894Rfs*2/+}$;*Chat-eGFP*, and their *Celsr3*$^{+/+}$;*Chat-eGFP* littermate controls, sections were labelled with primary antibodies against GFP (chicken anti-GFP, 1:1000, Aves Labs GFP-1020, RRID:AB_10000240) and PV (guinea pig anti-PV 1:1500, Swant PVG-213, RRID:AB_2650496) for two nights at 4℃. Sections were then labeled with secondary antibodies (donkey anti-chicken Alexa Fluor 488, 1:200, Jackson IR 703-545-155, RRID:AB_2340375, donkey anti-guinea pig Alexa Fluor 647, 1:200, Jackson IR, 706-605-148, RRID:AB_2340476) for five hours at room temperature. In *Wwc1*$^{W88C/+}$ mice and their littermate controls, sections were labeled with primary antibodies against ChAT (goat anti-ChAT, 1:200, Millipore AB144P, RRID:AB_207975) and PV (as above) for two nights at 4℃. Sections were then labeled with secondary antibodies (donkey anti-goat Alexa Fluor 488, 1:200, ThermoFisher #A-11055, RRID:AB_2534102, and donkey anti-guinea pig Alexa Fluor 647, see above) for five hours at room temperature. A subset of sections was co-labelled for GFP and ChAT to ensure the GFP transgene was uniformly expressed in Chat+ interneurons and this showed ~100% colocalization. Images (z-stack, tile) were captured using a Zeiss LSM800 confocal microscope with a 20x 0.8NA NA Plan Apo objective. Interneurons were counted using Spots in Imaris (Bitplane), with 25 μm and 15 μm thresholds for CINs and PVINs, respectively. Interneuron density was calculated as #interneurons/volume and expressed as the number of interneurons per cubic millimeter.

Nissl staining was performed on 50 μm sections using 435/455 blue fluorescent Nissl stain (Invitrogen N21479). For Drd1a-tdTomato staining, 100 μm sections were stained with a primary antibody against RFP (rabbit anti-RFP, 1:1500, overnight 4℃, Rockland 600-401-379, RRID:AB_2209751) and a secondary antibody (donkey anti-rabbit Alexa Fluor™ 546, 1:1000, overnight 4℃, ThermoFisher A10040). Cortical layers were stained in 50 μm coronal sections using primary antibodies against transcription factors Ctip2 (rat anti-Ctip2, 1:1000, Abcam ab18465, RRID:AB_2064130), Foxp2 (rabbit anti-Foxp2, 1:1000, Abcam ab16046, RRID:AB_2107107), and Satb2 (mouse anti-Satb2, 1:50, Abcam ab51502, RRID:AB_882455) for 2 nights at 4℃, followed by incubation overnight at 4℃ in secondary antibodies (goat anti-rat Alexa Fluor™ 488, goat anti-rabbit Alexa Fluor™ 647, goat anti-mouse Alexa Fluor™ 546, all 1:1000, ThermoFisher A-11030, A-21245, A-11006, RRID:AB_2534089, RRID:AB_141775, RRID:AB_2534074). Nissl and Drd1a-tdTomato images were assessed qualitatively. Cortical layer image volumes were imported into Imaris (Bitplane). Cortical layer thickness was measured manually, and normalized to total cortical thickness. All graphing and statistical analysis was done in Prism (GraphPad).

**3D pose analysis: data acquisition, processing, and modeling**: Mice with a median age of eight weeks were tested. Analysis was performed using tools and procedures provided by the Datta Lab, and following previous publications (*23*). The following programs and versions were used: kinect2-nidaq (v0.2.4-alpha), moseq2-extract (v1.1.2), moseq2-pca (v1.1.3), moseq2-model (v1.1.2), moseq2-viz (v1.2.0). First, we used the program kinect2-nidaq (v0.2.4-alpha) to collect raw depth frames from a Microsoft Kinect v2 device, mounted above the arena. Mice were placed at the bottom edge of a black polyethylene bucket measuring 43 cm in diameter and 35 cm in height (Tamco Industries, Cat #14317) and allowed to freely explore the arena for 20 minutes. Frames were collected at 30 Hz, and each frame was composed of 512 x 424 pixels, with each pixel containing a 16-bit unsigned integer specifying the distance of that pixel (in millimeters) from the sensor. After each session, frames were gzip compressed and moved to another computer for offline analysis.

The raw data for each recording session was extracted using the program moseq2-extract (v1.1.2), largely using the default parameters and the flip model "flip_classifier_k2_c57_10to13weeks.pkl" supplied by the Datta Lab. Briefly, the mouse's center and orientation were found using an ellipse fit on the pixels identified as "mouse". Then, an 80x80 pixel box was drawn around the mouse, and the mouse was rotated to face the right-hand side. All extraction results were assessed for quality of extraction by a human watching a movie visualization of the data and also by comparing the distributions of height, width, length, and area, following best practices as described by the Datta Lab. To account for variation in the depth images not due to changes in pose dynamics, extracted data were passed through a denoising convolutional autoencoder, as previously described (2).

Next, we used the program moseq2-pca (v1.1.3) to project the extracted depth frames onto the first 10 learned principal components (PCs), forming a 10-dimensional time series that described the mouse's 3D pose trajectory. Quality of the PCA model was assessed by examining a visualization of the pixel weights assigned by each principal component as well as the cumulative distribution of the percent variance explained by each principal component. This program was also used to generate a model-free changepoint analysis, which describes an empirical syllable duration distribution, found without any model constraints.

Then we used the program moseq2-model (v1.1.2) and the 10-D PCA-transformed data to train a series of autoregressive hidden Markov models (AR-HMM, a.k.a "moseq model"). Each state was described by a vector autoregressive process that captures the evolution of the 10 PCs over time and a hidden markov model that captures the switching dynamics between these states. For all models, we used the following parameters: "--max-states 100 --robust". We determined the best value for the hyperparameter kappa, which affects the timescale of discovered behavioral syllables. For this we trained a family of 100 models for 200 iterations each, with kappa values ranging logarithmically from 100,000 to 1,000,000,000. The best kappa parameter was determined by minimizing the absolute difference in the mean syllable duration between a given AR-HMM model fit and the mean block duration found via a model-free changepoint analysis (see above). For the results presented in this manuscript, we found a kappa value of 31,992,671 satisfied these criteria. Then, we trained a family of 100 models for 1,000 iterations each, using this discovered optimal kappa value. To choose an appropriate model from this family, we examined the aggregate log-likelihood value for each model and chose the model with the median log-likelihood to carry forward to downstream analysis.

**3D pose analysis: Behavioral usage and transition matrix analysis**: Syllable usage was calculated by counting the number of occurrences of each syllable and dividing by the total sum of all syllable occurrences within a recording session, converting syllable usage into a percentage. The number of syllables analyzed were cutoff based on the global usage across all sessions, eliminating syllables which were not performed by any animals in the study. Transition matrices were calculated by counting the total number of occurrences where syllable *A* transitions into syllable *B* (for all syllables) and normalizing by the sum of the matrix (bigram normalization). Statistical testing for syllable usage follows the previously published procedures. Briefly, for each group comparison of interest and each syllable, we took 1,000 bootstrap samples (sampling with replacement within a group) of the given syllable's usage for each group and performed a z-test on these two distributions. Finally, we use the Benjamini-Hochberg procedure (statsmodels.stats.multitest.multipletests) to correct for multiple hypothesis testing and control the false discovery rate.

**3D pose analysis: Entropy and Entropy Rate Analysis**: Entropy and entropy rate was calculated using standard formulas:

$$Entropy(Y) = -\sum_{i} \mu_i log_2(\mu_i)$$

$$EntropyRate(Y) = -\sum_{ij} \mu_i P_{ij}\, log_2(P_{ij})$$

Where $Y$ is an observed syllable emission sequence, $\mu$ is the asymptotic distribution of the markov chain, approximated by empirical normalized usage emissions, and $P$ is an empirical bigram normalized transition matrix. To enable global comparison between controls and mutants irrespective of mouse line, each animal's entropy and entropy rate was normalized by the corresponding median value from the respective sex and line matched control sample. Statistical testing was performed using a Mann Whitney U test.

**3D pose analysis: Behavioral Linear Discriminant Analysis**: Linear Discriminant Analysis (LDA) was performed using the scikit-learn implementation using the eigen solver and 2 components. Individual normalized usage or bigram transition probabilities were fed as input to the LDA model, including group labels. This data was split into train and validation sets in a 70:30 stratified ratio. We searched over the hyperparameter "shrinkage" using a 5-fold stratified cross-validation approach using only the train subset and found little effect on the model performance against the never before-seen validation set. Final models were trained using the entire train set, and evaluated on the never-before-seen validation set. We also performed a permutation test (sklearn.model_selection.permutation_test_score), wherein we train a family of models against 100 sets of randomly permuted labels, and compare the distribution of model scores against shuffled data vs our final model and calculate a p-value. Results were plotted with seaborn and matplotlib.

**3D pose analysis: Cosine Distances**: We first calculated the cosine distance of every animal to every other animal in the dataset using normalized usage emissions and the scipy function `pdist` with the parameter `metric="cosine"`. Then for each animal, we computed the mean distance to all other animals either A) within the same group as the current animal (within-group) or B) outside of the same group as the current animal (between-group). Data was then plotted using seaborn and matplotlib, showing the mean and 95% CI of the within- and between-group distance distributions observed for each group of animals.

**Statistical Analyses:** Statistical analysis of all behavior paradigms with the exception of Motion Sequencing was completed using Graphpad Prism (Graphpad Software Inc., La Jolla, CA). Drug rescues were performed using R. Prepulse inhibition (PPI) of the acoustic startle reflex was analyzed using two-way ANOVA across all three prepulses. Spearman's correlation coefficient was performed to determine if there was a correlation between startle amplitude and prepulse inhibition. Rotarod was analyzed across all nine trials using two-way ANOVA with repeated measures. Marble burying was analyzed using the Mann Whitney test. Time spent grooming, time/grooming bout, and then number of grooming events were analyzed using Mann-Whitney Test. Open field distance traveled and rearing events were analyzed using two-way ANOVA with repeated measures and Bonferroni multiple correction testing. Unless otherwise noted, two-way ANOVA results are presented as the main effect of genotype throughout the text. For reinforcement learning, Mann-Whitney tests were used to compare time to completion, and t-tests with Welch's correction and Holm-Šídák's multiple comparison correction were used to compare rewards obtained. For analyzing time spent to earn 30 rewards over four days, or latency to retrieve first reward, ANOVA using ordinary least-squares algorithm to fit a linear model was implemented, using day and genotype as input factors. Statistical analyses of drug treatment effects on PPI and rearing were performed using multiple linear regression (Type III sum of squares) and by fitting a linear equation to the input and output variables. For PPI, the input variables (i.e., factors) were decibel settings (factor levels 71db, 77db, 81db), genotype (wildtype and mutant), and drug treatment (vehicle or aripiprazole). The outcome variable was PPI. For rearing, the input factors were time (0-10 min, 10-20 min, 20-30 min), genotype (wildtype and mutant), and drug treatment (vehicle and aripiprazole). The outcome variable was rearing counts. A detailed explanation of the statistical analyses for Motion Sequencing can be found in the supplemental methods. Briefly, for behavioral usage Mann Whitney U analysis was performed followed by Benjamini-Hochberg multiple comparisons correction. LDA analysis was completed using scikit-learn implementation.

### A Celsr3

S1894Rfs*2    C1906Y

```
Homo sapiens          GSELQGLKVKQLHVGGLPPGSAEEAPQGLVGCIQGVWLGSTPSGSPALLPPSHRVNAEPG 1943
Mus musculus          GGELQGLKVKQLHVGGLPPSSKEEGHQGLVGCIQGVWIGFTPFGSSALLPPSHRVNVEPG 1934
Rattus norvegicus     GSELEGLKVKHLHVGGPPPSSKEEGPQGLVGCIQGVWTGFTPFGSSALPPPSHRINVEPG 1934
Macaca mulatta        GSELQGLKVKQLHVGGLPPGSAEEAPQGLVGCIQGVWLGSTPSGSPALLPPSHQVNAEPG 1943
Macaca fascicularis   GSELQGLKVKQLHVGGLPPGSAEEAPQGLVGCIQGVWLGSTPSGSPALLPPSHQVNAEPG 1943
Callithrix jacchus    GSELQGLKVKQLHVGGLPPGSAEEAPQGLVGCIQGVWLGSTPSGSPALLPPSHGVNVEPG 2064
Cricetulus griseus    GGELQGLKVKQLHVGGLPPTSKEEGPQGLVGCIQGVWIGYTPFGSSALPPPSHRVNVEPG 1986
Cavia porcellus       GTELHGLQVKQLHVGGLPLSSKEEAPQGLVGCVQGVWLGSAPLGSPALLPPSHRVNVEPG 1933
Xenopus tropicalis    GSELHGLRVKNLYIGGVS--GPREVQNGFEGCIQGVRLGETPSGI-TLPKPSSALNVKPG 2270
Danio rerio           GNEINGVKVKHLHVGGVL--GSGEVQNGIRGCIQGVRLGVRP-DSPALPRPSRTIKVETG 2314
                      * *:.*::**:*::**    .  *  :*: **:***  *  * .  :*  **   ::.: * (aa)
```

### B Wwc1

W88C

```
Homo sapiens          EAYDPQVGDYFIDHNTKTTQIEDPRVQWRREQEHMLKDYLVVAQEALSAQKEIYQVKQQR 120
Mus musculus          EAYDPQVGDYFIDHNTKTTQIEDPRVQWRREQEHMLKDYLVVAQEALSAQKEIYQVKQQR 120
Rattus norvegicus     EAYDPQVGDYFIDHNTKTTQIEDPRVQWRREQEHMLKDYLVVAQEALSAQKEIYQVKQQR 120
Macaca mulatta        EAYDPQVGDYFIDHNTKTTQIEDPRVQWRREQEHMLKDYLVVAQEALSAQKEIYQVKQQR 120
Macaca fascicularis   EAYDPQVGDYFIDHNTKTTQIEDPRVQWRREQEHMLKDYLVVAQEALSAQKEIYQVKQQR 120
Callithrix jacchus    EAYDPQVGDYFIDHNTKTTQIEDPRVQWRREQEHMLKDYLVVAQEALSAQKEIYQVKQQR 120
Cricetulus griseus    EAYDPQVGDYFIDHNTKTTQIEDPRVQWRREQEHMLKDYLVVAQEALSAQKEIYQVKQQR 120
Cavia porcellus       EAYDPQVGDYFIDHNTKTTQIEDPRVQWRREQEHMLKDYLVVAQEALSAQKEIYQVKQQR 120
Xenopus tropicalis    ESYDTQVGVYYIDHNSQTTQIEDPRVQWRREQERMLKDYLVLAQEALLAQKEIYQVKQQR 120
Danio rerio           EAYDPHVGAYYVDHNTKSTQLEDPRAQWQREQELMLHDYLNVVQEALSAQKEIYQVKEQR 120
                      *:** :** *::***:::**:****.**:**** **:*** :.**** *********:** (aa)
```

Figure S1 - Nasello et al. (2023)

**Figure S1: *Celsr3* and *Wwc1* amino acid sequence in the region of the TD associated mutations.** Arrows indicate location of change in amino acid sequences. **(A)** C1906Y and S1894Rfs*2 are in a conserved region of Celsr3. **(B)** W88C is in a conserved region of Wwc1.

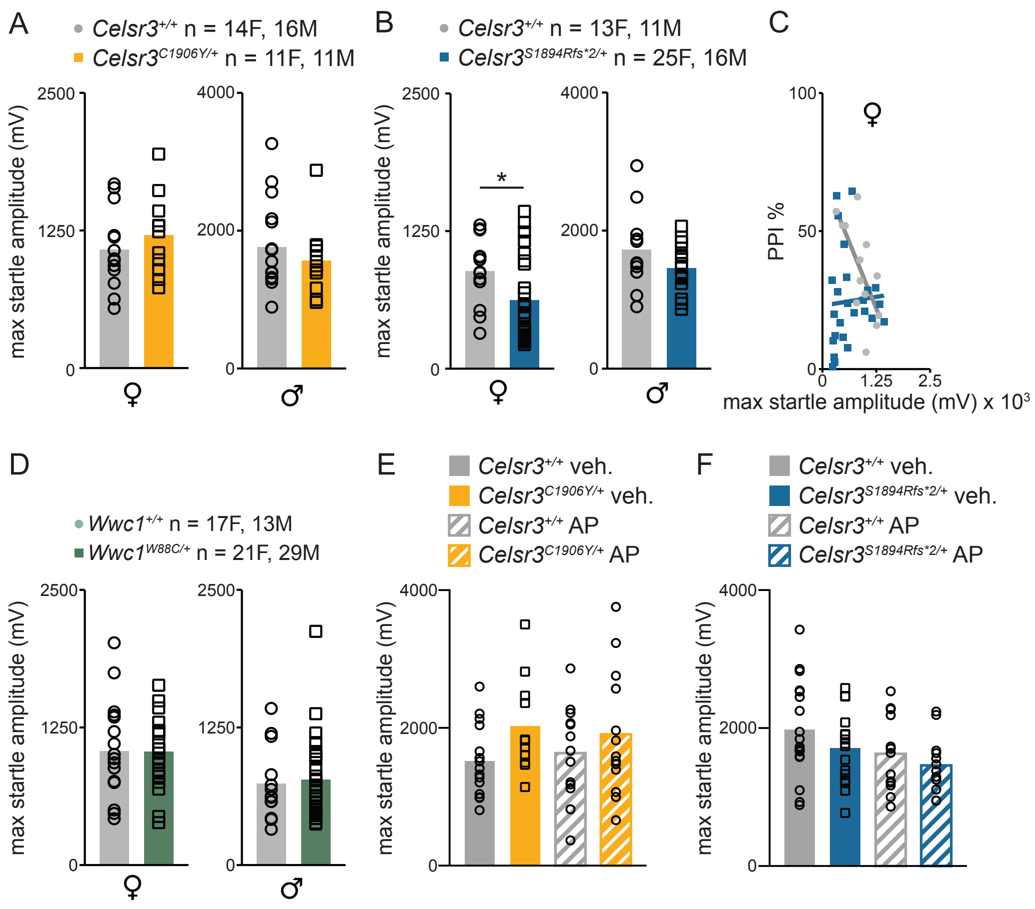

Figure S2 - Nasello et al. (2023)

**Figure S2: Startle amplitude is unaffected except for *Celsr3*$^{S1894Rfs*2/+}$ females.** Startle responses across multiple trials of 120 dB intensity were recorded through an accelerometer located underneath each mouse. **(A)** *Celsr3*$^{C1906Y/+}$ mice had comparable startle responses to controls (Female: *p=0.798* and male: *p=0.403*). **(B)** Female *Celsr3*$^{S1894Rfs*2/+}$ mice had a significant reduction in startle amplitude (*p=0.038*). Male *Celsr3*$^{S1894Rfs*2/+}$ mice had decreased startle amplitude that did not reach statistical significance (*p=0.146*). **(C)** *Celsr3*$^{+/+}$ females had a correlation between increased startle amplitude and decreased PPI (female: r=-0.648 and *p=0.019*) while female *Celsr3*$^{S1894Rfs*2/+}$ had no correlation between startle amplitude and levels of PPI (r=0.267 and *p=0.20*). **(D)** Female and male *Wwc1*$^{W88C/+}$ mice had comparable startle responses to controls (female: *p=0.9567* and male: *p=0.9131*). **(E, F)** Startle responses in mice treated with either saline vehicle or aripiprazole. **(A, B, D)** Unpaired t-test. **(C)** Spearman's correlation coefficient. **(E, F)** One-way ANOVA. *Celsr3*$^{+/+}$/*Celsr3*$^{C1906Y/+}$ female (n=14/11); *Celsr3*$^{+/+}$/*Celsr3*$^{C1906Y/+}$ male (n=16/11); *Celsr3*$^{+/+}$/*Celsr3*$^{S1894Rfs*2/+}$ female (n=13/25); *Celsr3*$^{+/+}$/*Celsr3*$^{S1894Rfs*2/+}$ male (n=11/29); *Wwc1*$^{+/+}$/*Wwc1*$^{W88C/+}$ female (n=17/20), *Wwc1*$^{+/+}$/*Wwc1*$^{W88C/+}$ male (n=13/29).

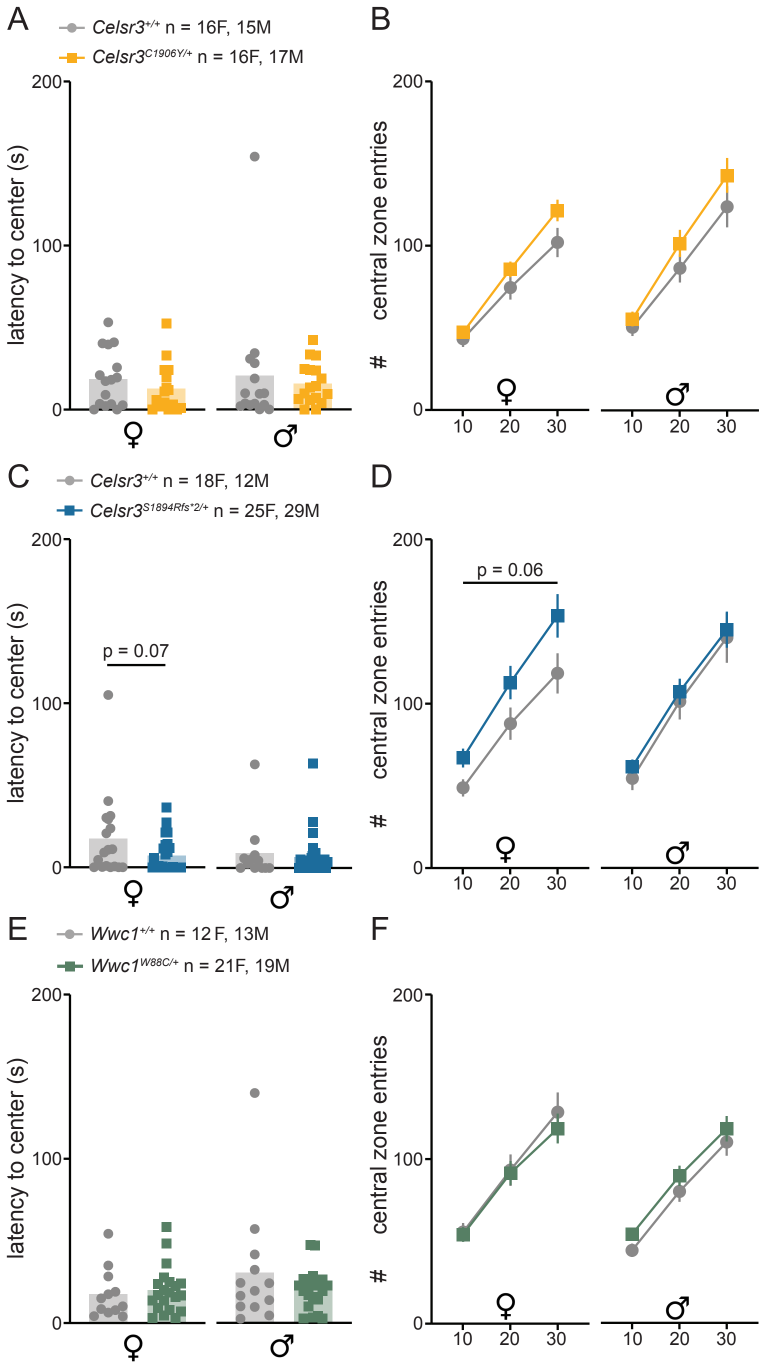

Figure S3 - Nasello et al. (2023)

**Figure S3: Latency to enter the center of the open field arena is normal but female *Celsr3*$^{C1906Y/+}$ and *Celsr3*$^{S1894Rfs*2/+}$ mice show slightly more central zone crossings. (A)** *Celsr3*$^{C1906Y/+}$ mice show similar latency to enter the central zone compared to controls (female: *p=0.61*, male: *p=0.59*). **(B)** Female *Celsr3*$^{C1906Y/+}$ mice made more central zone entries over time [Time x genotype interaction: F(2, 60)=5.69, *p=0.005*] but genotype as main effect was not significant [F(1, 30)=1.88, *p=0.18*]. Differences became more apparent as testing progressed (30 minute: *p=0.09*). The number of central zone entries was normal for male *Celsr3*$^{C1906Y/+}$ mice [time x genotype interaction: F(2, 60)=2.04, *p=0.139* and main effect genotype F(1, 60)=1.3, *p=0.266*]. **(C)** *Celsr3*$^{S1894Rfs*2/+}$ female mice showed a trend-level effect for reduced latency to enter the central zone compared to controls, while *Celsr3*$^{S1894Rfs*2/+}$ male mice showed similar latency compared to controls (female: *p=0.0721*, male: *p=0.6672*). **(D)** Female *Celsr3*$^{S1894Rfs*2/+}$ mice show a slight increase in the number of central zone entries [time x genotype interaction F(2, 82)=2.17, *p=0.121*, main effect genotype: F(1, 41)=3.61, *p=0.065*]. Significance was nearly reached at the end of the testing period (30 minute: *p=0.052*). The number of central zone entries was normal for male *Celsr3*$^{S1894Rfs*2/+}$ mice [time x genotype F(2, 78)=0.35, *p=0.965* and genotype main effect: F(1,39)=0.1922, *p=0.65*]. **(E)** *Wwc1*$^{W88C/+}$ mice have a similar latency to enter the center of the arena compared to controls (female *p=0.66*, male: *p=0.74*). **(F)** The number of central zone entries was normal for female and male *Wwc1*$^{W88C/+}$ mice. [Female: time x genotype: F(2, 64)=1.6, *p=0.21*, genotype main effect: F(1, 32)=0.15, *p=0.703*, Male: time x genotype: F(2, 58)=0.56, *p=0.95*, F(1,29)=1.1, *p=0.29*]. **(A, C, E)** Unpaired t-test **(B, D, F)** Two-way RM-ANOVA, Bonferroni's multiple comparisons test. *Celsr3*$^{+/+}$/*Celsr3*$^{C1906Y/+}$ female (n=16/16); *Celsr3*$^{+/+}$/*Celsr3*$^{C1906Y/+}$ male (n=15/17); *Celsr3*$^{+/+}$/*Celsr3*$^{S1894Rfs*2/+}$ female (n=18/25); *Celsr3*$^{+/+}$/*Celsr3*$^{S1894Rfs*2/+}$ male (n=12/29); *Wwc1*$^{+/+}$/*Wwc1*$^{W88C/+}$ female (n=12/21), *Wwc1*$^{+/+}$/*Wwc1*$^{W88C/+}$ male (n=13/18).

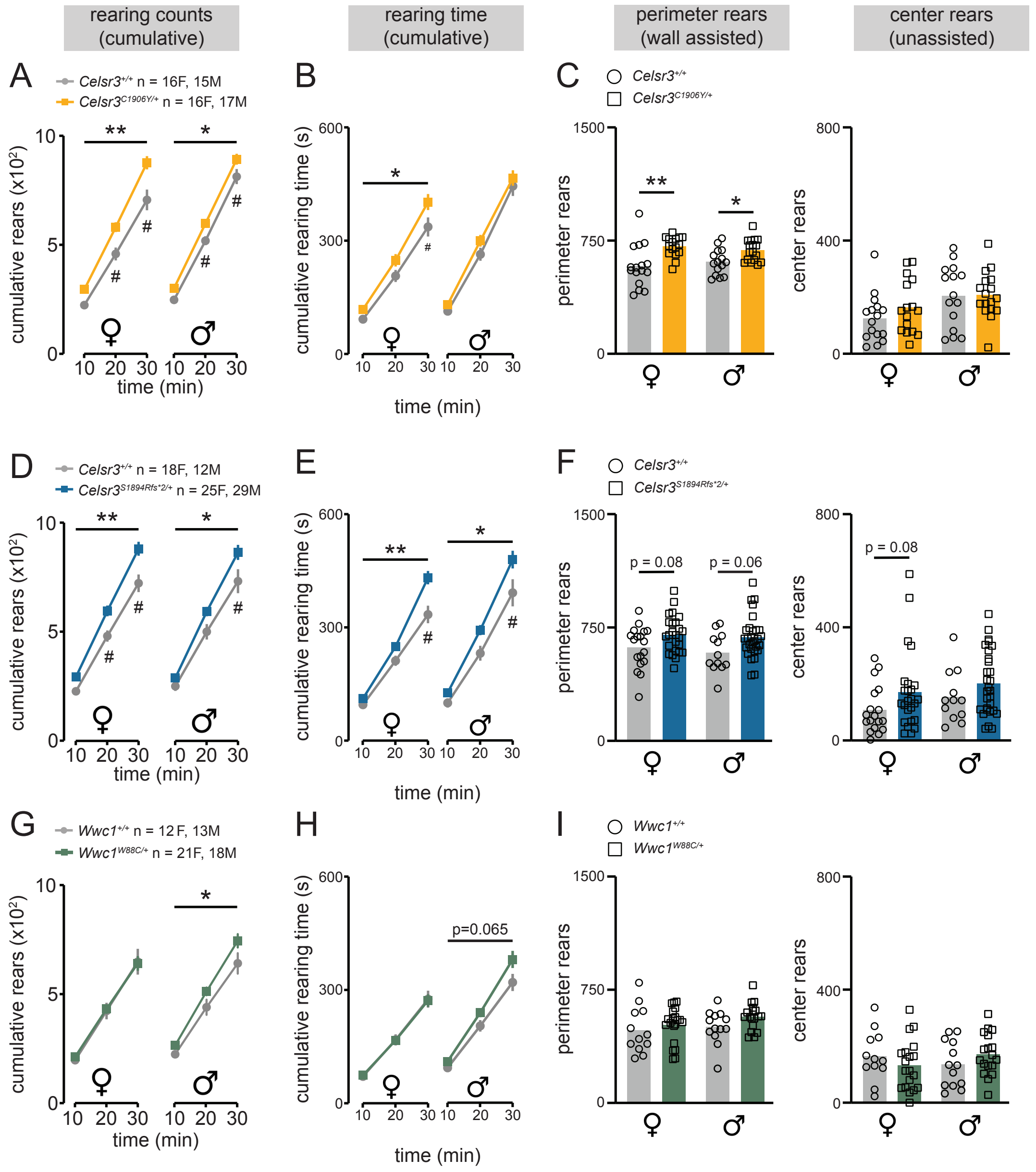

Figure S4 - Nasello et al. (2023)

**Figure S4: Rearing behavior in TD mouse models is increased. (A, D, G)** Cumulative rearing events. **(B, E, H)** Cumulative time spent rearing by each ten-minute interval. **(C, F, I)** Endpoint rearing counts performed along the perimeter (left) vs. the center (right) of the open field arena. **(A)** Total number of rearing events are increased for *Celsr3*$^{C1906Y/+}$ mice [Female: time x genotype interaction $F(2, 60)=5.99$, *p=0.004*, main effect genotype: $F(1, 30)=10.77$, *p=0.003*] [Male: time x genotype interaction $F(2, 60)=0.72$, *p=0.49*, main effect genotype: $F(1, 30)=7.44$, *p=0.011*]. Significance increased as time progressed (Female: 0-10, *p=0.24*; 0-20, *p=0.0095*; 0-30, *p=0.0002*, Male: 0-10, *p=0.24*; 0-20, *p=0.028*; 0-30, *p=0.026*). **(B)** Total rearing time was increased for *Celsr3*$^{C1906Y/+}$ mice. [Female: time (x) genotype interaction $F(2, 60)=2.31$, *p=0.11*, main effect genotype: $F(1, 30)=4.3$, *p=0.045*; Male: time (x) genotype interaction $F(2, 60)=3.9$ *p=0.024*, main effect genotype $F(1, 30)=0.45$, p=0.52]. **(C)** *Celsr3*$^{C1906Y/+}$ mice show more rearing events along the perimeter of the arena (female: *p=0.0010*, male: *p=0.023*) but not within the central zone (female: *p=0.25*, male: *p=0.85*). **(D)** Total number of rearing events are increased for *Celsr3*$^{S1894Rfs*2/+}$ females [Females: time x genotype interaction $F(2, 82)=5.8$, *p=0.004*, main effect genotype: $F(1, 41)=10.8$, *p=0.002*] and significance increased as time progressed (0-10, *p=0.25*; 0-20, *p=0.0084*; 0-30, *p=0.0002*). *Celsr3*$^{S1894Rfs*2/+}$ males also rear more [Males: time x genotype interaction $F(2, 78)=3.6$, *p=0.03*, main effect genotype: $F(1, 39)=4.6$, *p=0.038*] and significance increased as time progressed (0-10, *p>0.99*; 0-20, *p=0.13*; 0-30, *p=0.013*). **(E)** Cumulative rearing time was increased for *Celsr3*$^{S1894Rfs*2/+}$ mice [Female: time X genotype interaction $F(2, 82)=12.2$, *p<0.0001*, main effect genotype: $F(1, 41)=8.1$, *p=0.007*] [Male: time x genotype interaction $F(2, 78)=2.9$, *p=0.059,* main effect genotype: $F(1, 39)=4.9$, *p=0.03*]. The greatest differences occurred during the last 10-minute time block (Female: 10, *p>0.999*, 20, *p=0.203*, 30, *p<0.0001*, Male: 10, *p>0.999*, 20, *p=0.130*, 30, *p=0.012*). **(F)** Rearing events along the perimeter (*p=0.08*) and within the central zone (*p=0.08*) of the arena trended upward for female *Celsr3*$^{S1894Rfs*2/+}$ mice. Rearing events along the perimeter (*p=0.06*) also trended upward for *Celsr3*$^{S1894Rfs*2/+}$ males, but not in the central zone (*p=0.24*). **(G)** Female *Wwc1*$^{W88C/+}$ mice reared a similar number of times as controls [time x genotype interaction $F(2, 64)=0.02$, *p=0.98*, main effect genotype: $F(1, 32)=0.35$, *p=0.56*]. Cumulative number of rearing events were increased for males (time x genotype interaction $F(2, 58)=2.2$, *p=0.12*, main effect genotype: $F(1, 29)=4.2$, *p=0.048*]. **(H)** Cumulative rearing time was similar for *Wwc1*$^{W88C/+}$ females compared to controls [time x genotype interaction $F(2, 64)=0.09$, *p=0.9*, main effect genotype: $F(1, 32)=0.0001$, *p=0.97*]. *Wwc1*$^{W88C/+}$ males showed a trend-level effect [time x genotype interaction $F(2, 58)=12.5$, *p=0.083*, main effect genotype: $F(1, 29)=3.6$, *p=0.065*]. **(I)** Female and male *Wwc1*$^{W88C/+}$ showed no changes to rearing events in either the perimeter (Female: *p=0.45*, Male: *p=0.12*) or central zone (Female: *p=0.341*, male: *p=0.24*). **(A,**

**B, D, E, G, H)** Two-Way RM ANOVA. **(C, F, I)** Mann Whitney Test. *Celsr3*$^{+/+}$/*Celsr3*$^{C1906Y/+}$ female (n=16/16); *Celsr3*$^{+/+}$/*Celsr3*$^{C1906Y/+}$ male (n=15/17); *Celsr3*$^{+/+}$/*Celsr3*$^{S1894Rfs*2/+}$ female (n=18/25); *Celsr3*$^{+/+}$/*Celsr3*$^{S1894Rfs*2/+}$ male (n=12/29); *Wwc1*$^{+/+}$/*Wwc1*$^{W88C/+}$ female (n=13/21), *Wwc1*$^{+/+}$/*Wwc1*$^{W88C/+}$ male (n=13/18).

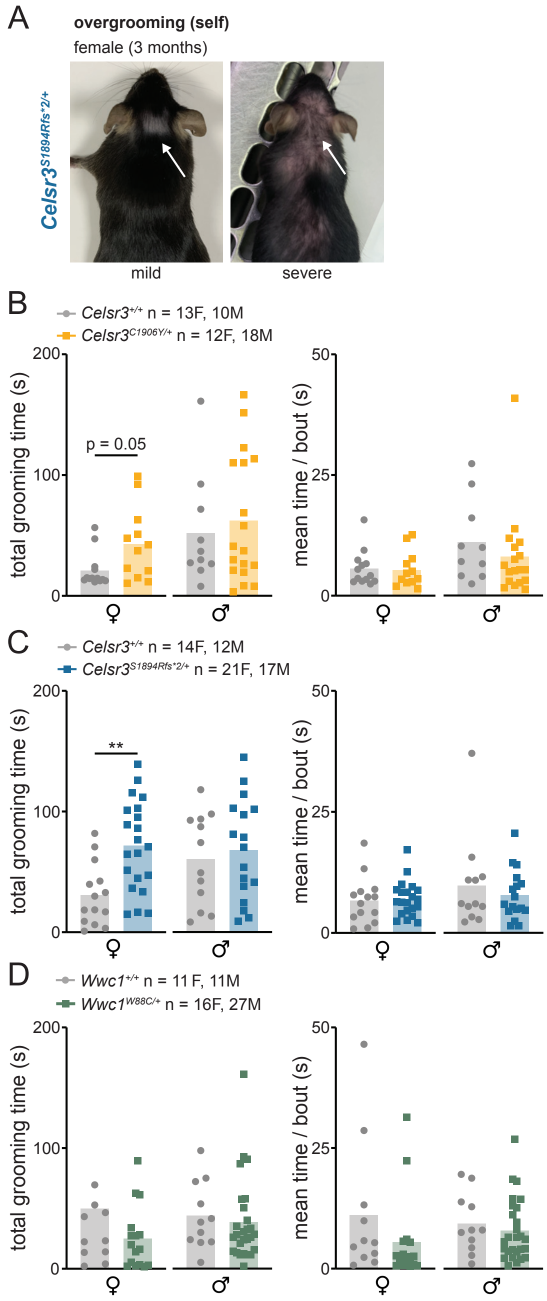

Figure S5 - Nasello et al. (2023)

**Figure S5: *Celsr3* TD mouse models spend more time grooming with normal bout duration.** **(A)** Representative photos of two over-groomed *Celsr3*$^{S1894Rfs*2/+}$ females, mild (left), severe (right). **(B)** Total time spent grooming was increased for *Celsr3*$^{C1906Y/+}$ females versus controls, but *Celsr3*$^{C1906Y/+}$ males did not show a difference (female: *p=0.05*, male: *p=0.87*). The mean time spent grooming per bout was normal for *Celsr3*$^{C1906Y/+}$ mice (female: *p=0.65*, male: *p=0.245*). **(C)** *Celsr3*$^{S1894Rfs*2/+}$ females spent more time grooming during the ten-minute testing period versus controls. *Celsr3*$^{S1894Rfs*2/+}$ males showed no differences versus controls (female: *p=0.001*, male: *p=0.62*). The mean time spent grooming per bout was normal for *Celsr3*$^{S1894Rfs*2/+}$ mice (female: *p=0.61*, male: *p=0.59*). **(D)** *Wwc1*$^{W88C/+}$ mice showed no changes to time spent grooming during the testing period (female: *p=0.3420*, male: *p=0.41*). Female *Wwc1*$^{W88C/+}$ mice displayed a trend towards a shorter mean time spent per grooming bout (female: *p=0.11*, male: *p=0.356*). **(A-D)** Mann Whitney Test. *Celsr3*$^{+/+}$/*Celsr3*$^{C1906Y/+}$ female (n=13/12); *Celsr3*$^{+/+}$/*Celsr3*$^{C1906Y/+}$ male (n=10/18); *Celsr3*$^{+/+}$/*Celsr3*$^{S1894Rfs*2/+}$ female (n=14/21); *Celsr3*$^{+/+}$/*Celsr3*$^{S1894Rfs*2/+}$ male (n=12/27); *Wwc1*$^{+/+}$/*Wwc1*$^{W88C/+}$ female (n=11/26), *Wwc1*$^{+/+}$/*Wwc1*$^{W88C/+}$ male (n=11/27).

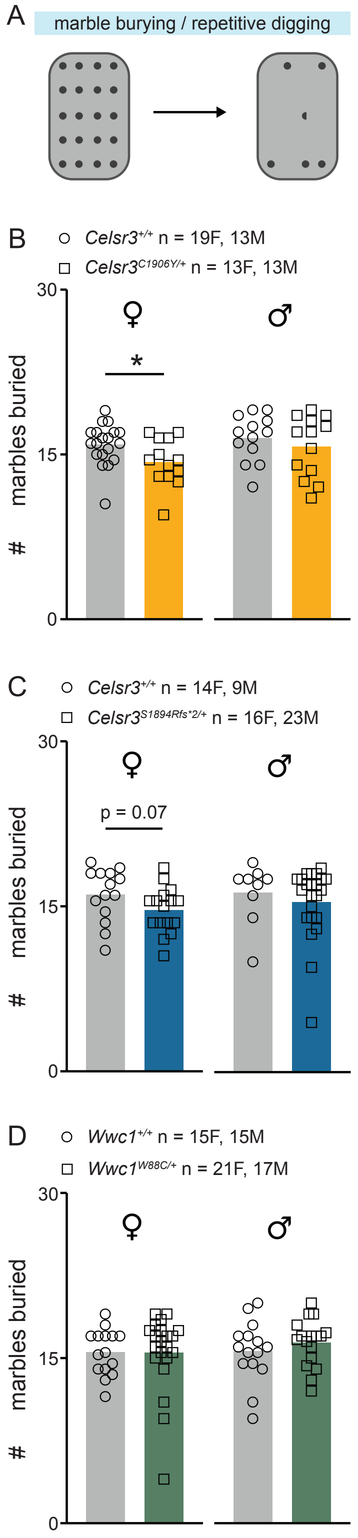

Figure S6 - Nasello et al. (2023)

**Figure S6: Female Celsr3 TD mice show less repetitive digging in a marble burying assay. (A)** Mice were placed in a standard polycarbonate box with a lattice of 4 X 5 marbles and 5 cm of bedding for 20 minutes. Any marble more than ⅔ buried was scored. **(B)** Female *Celsr3*$^{C1906Y/+}$ mice bury less marbles versus controls (*p=0.032*). No differences are detected for *Celsr3*$^{C1906Y/+}$ males versus controls (*p=0.6*). **(C)** *Celsr3*$^{S1894Rfs*2/+}$ females bury slightly less marbles versus controls, trending towards significance (*p=0.07*) whereas *Celsr3*$^{S1894Rfs*2/+}$ males bury equivalent numbers versus controls (*p=0.389*). **(D)** *Wwc1*$^{W88C/+}$ mice bury equivalent numbers of marbles compared to controls (Female: *p=0.383*, male: *p=0.438*). **(A-D)** Mann Whitney Test. *Celsr3*$^{+/+}$/*Celsr3*$^{C1906Y/+}$ female (n=19/13); *Celsr3*$^{+/+}$/*Celsr3*$^{C1906Y/+}$ male (n=13/13); *Celsr3*$^{+/+}$/*Celsr3*$^{S1894Rfs*2/+}$ female (n=14/16); *Celsr3*$^{+/+}$/*Celsr3*$^{S1894Rfs*2/+}$ male (n=9/23); *Wwc1*$^{+/+}$/*Wwc1*$^{W88C/+}$ female (n=15/21), *Wwc1*$^{+/+}$/*Wwc1*$^{W88C/+}$ male (n=15/17).

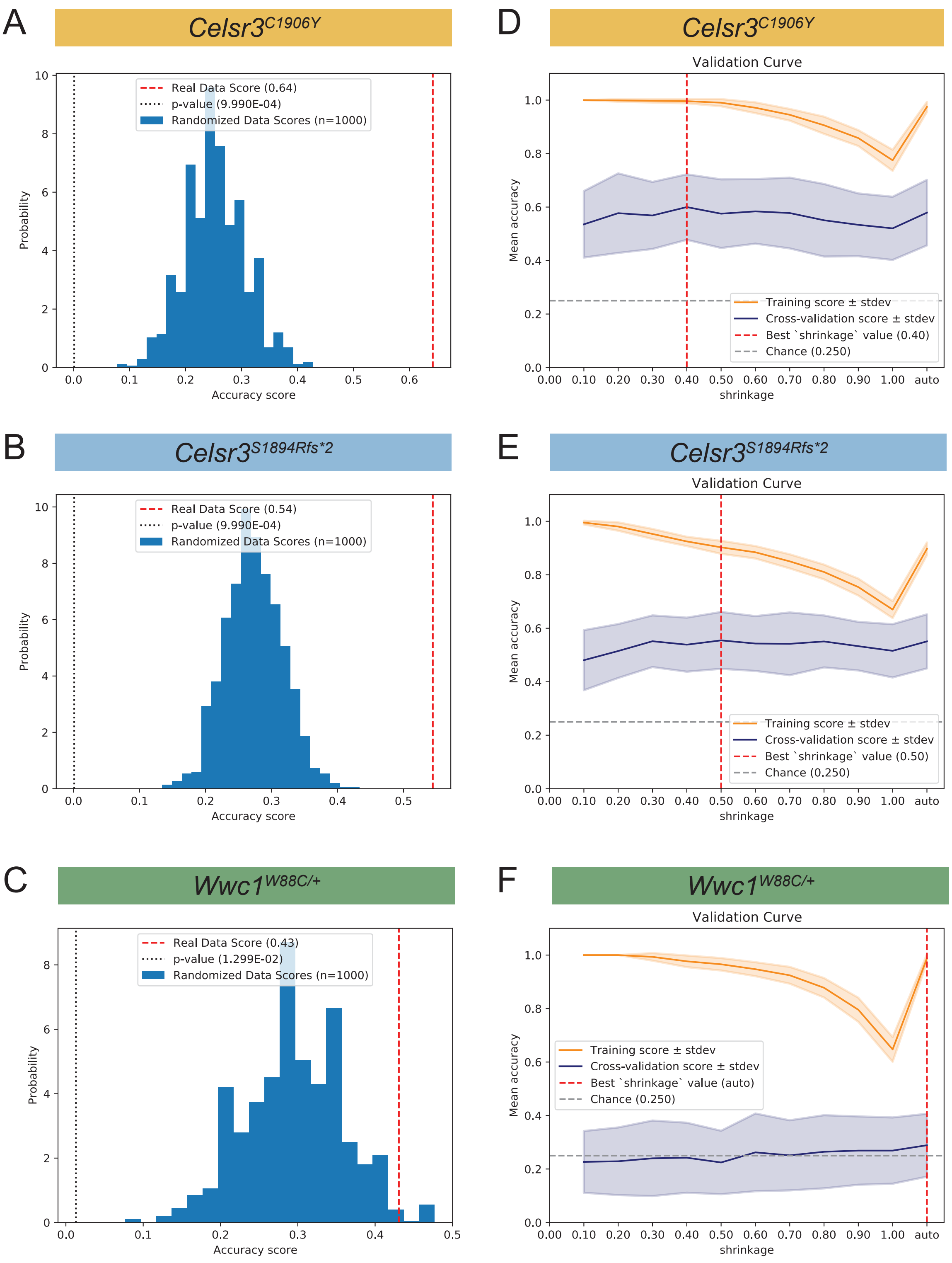

Figure S7 - Nasello et al. (2023)

**Figure S7: Classification of TD mice using linear discriminant analysis. (A, B, C)** Permutation plots corresponding to the LDA models in Fig. 4D for **(A)** *Celsr3*$^{C1906Y/+}$, **(B)** *Celsr3*$^{S1894Rfs*2/+}$ and **(C)** *Wwc1*$^{W88C/+}$. Histogram bars correspond to the overall classification accuracies from a family of LDA models trained using random permutations of group labels relative to input features. The overall accuracy of the real model is indicated by a red dashed line. The p-value, against the null hypothesis that features and group labels are independent, is indicated by the black dotted line. **(D-F)** Validation curves corresponding to the LDA models in Figure 4D for **(D)** *Celsr3*$^{C1906Y/+}$, **(E)** *Celsr3*$^{S1894Rfs*2/+}$, and **(F)** *Wwc1*$^{W88C/+}$.

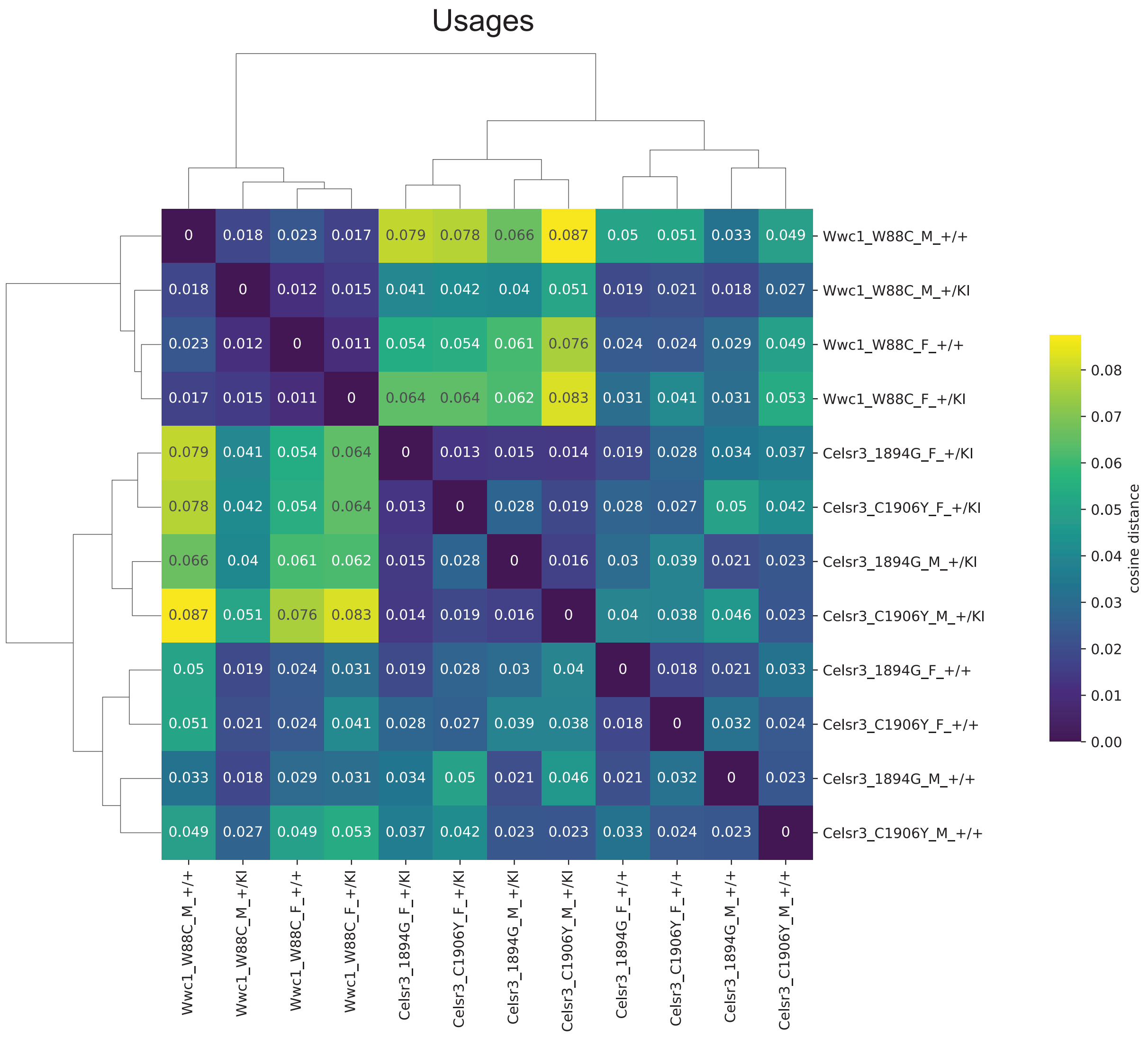

Figure S8 - Nasello et al. (2023)

**Figure S8: Classification of TD mice using cosine distances.** Heatmap showing cosine distances computed on mean usage frequencies between all group pairs. Dendrogram computed with linkage method “complete”. Plot showing the mean ± 95% CI of within-group (blue) or between-group (orange) cosine distances, computed on normalized usage frequencies, for each indicated group along the x-axis.

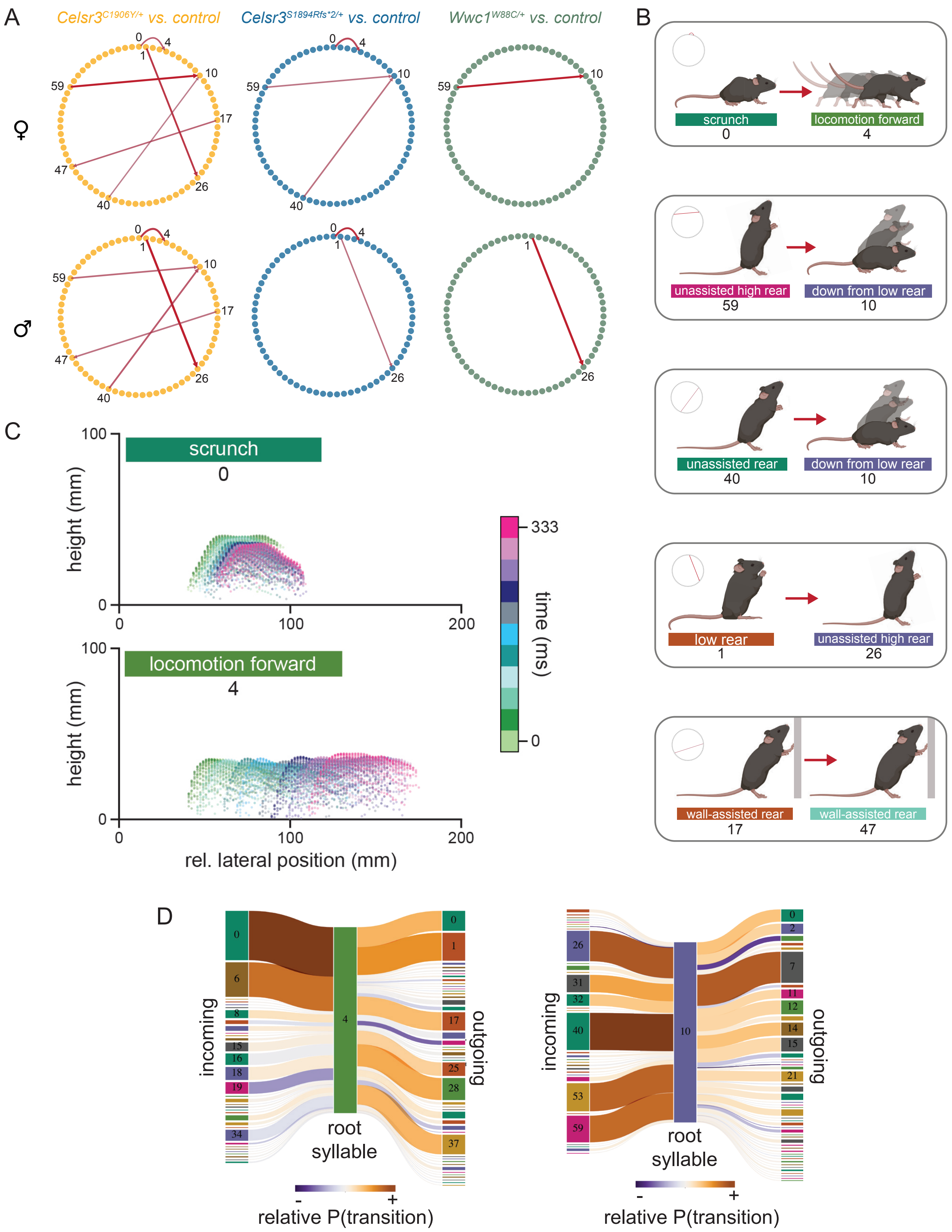

Figure S9 - Nasello et al. (2023)

**Figure S9: Commonly upregulated transitions between syllables in mutant mice. (A)** State maps pruned to show select "common" transitions shared across mutant mice. Circles represent syllables and red arrows show transitions between syllables that are upregulated in mutant relative to control mice. Numbers indicate syllable ID numbers/identifiers. **(B)** Illustrative examples for transitions between two syllables that have higher probability of occurring in mutant versus control mice. Illustrations were generated using BioRender.com. **(C)** Point clouds showing the three-dimensional pose data from two selected syllables, "scrunch" (0, top) and "locomotion forward" (4, bottom). Syllable duration is typically ~300 ms and color scale represents time. **(D)** Syllable flow diagrams showing example root syllables and the syllables that are most likely to precede them (incoming) and follow them (outgoing) in *Celsr3*$^{C1906Y/+}$ male mice relative to *Celsr3*$^{+/+}$ male mice.

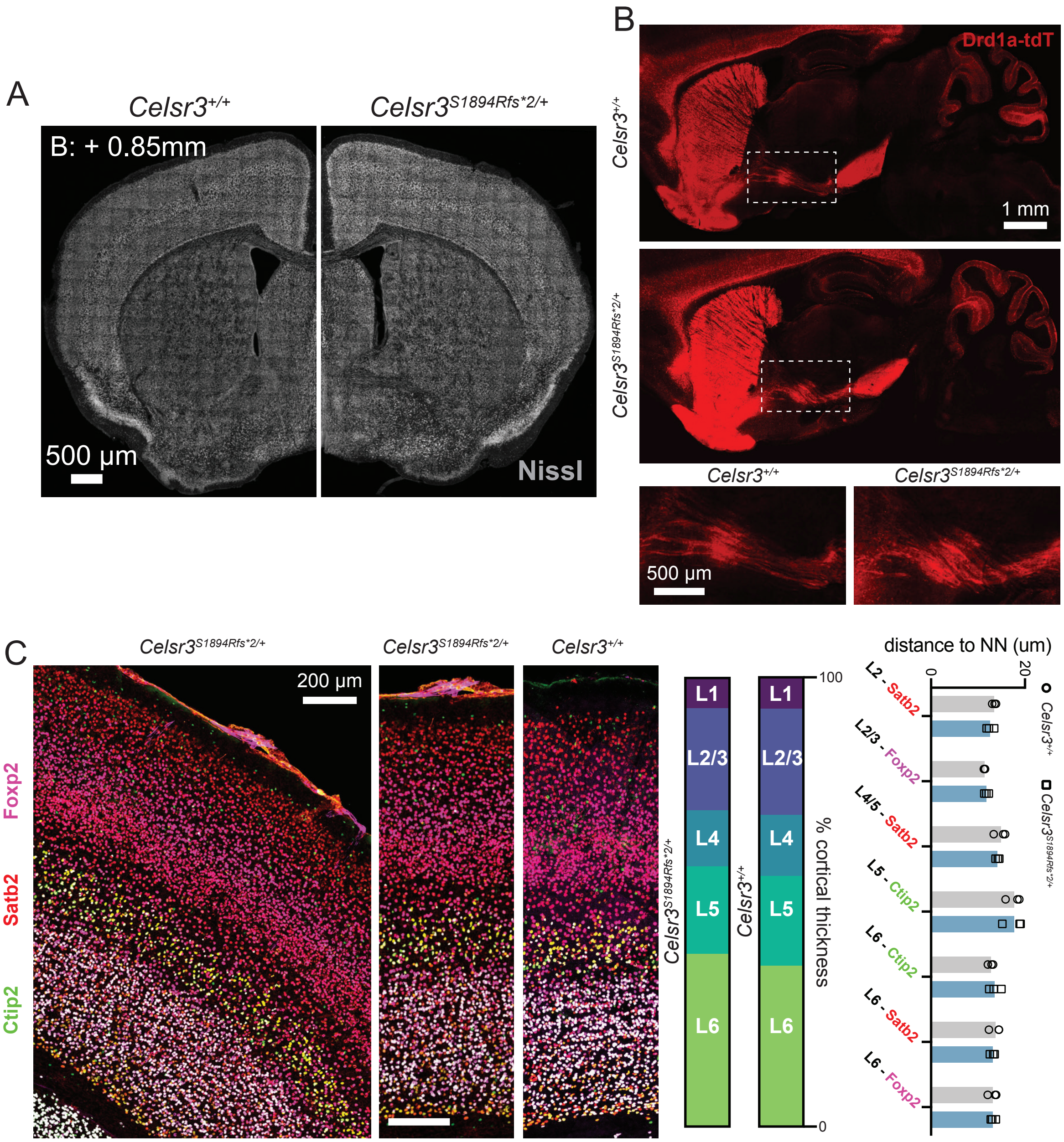

Figure S10 - Nasello et al. (2023)

**Figure S10: Gross neuroanatomical changes were not detected in *Celsr3*$^{S1894Rfs*2/+}$ mice. (A)** Coronal brain slices of *Celsr3*$^{+/+}$ (left) and *Celsr3*$^{C1906Y/+}$ mutant (right) mice labeled with Nissl show comparable neuronal density and patterning (*Celsr3*$^{+/+}$ n=3, *Celsr3*$^{S1894Rfs*2/+}$ n=2). **(B)** Sagittal brain slices of wild type (top) and mutant (bottom) mice both expressing *tdTomato* (tdT) in *Drd1a*+ (striatonigral) neurons are similar (*Celsr3*$^{+/+}$ n=3, *Celsr3*$^{S1894Rfs*2/+}$ n=3). Insets show fiber tracts. **(C)** Cortical layer markers Ctip2 (green), Satb2 (red), and Foxp2 (magenta) in coronal sections show normal laminar organization of somatosensory cortex (*Celsr3*$^{+/+}$ n=3, *Celsr3*$^{S1894Rfs*2/+}$ n=3). Relative (%) layer thicknesses (shown in parts-of-whole graphs) were similar, and nearest neighbor (NN) distances of populations within layers (right) were unchanged.

**Table S1 – Statistical testing of behavioral data.** Statistically significant ($p<0.05$) and trend level effects ($p<0.1$) values are highlighted in red.

| | | Test group | | | | | |
|---|---|---|---|---|---|---|---|
| **Behavior assay** | **Statistical test** | ***Celsr3^+/+^/ Celsr3^C1906Y/+^*** **Female** | ***Celsr3^+/+^/ Celsr3^C1906Y/+^*** **Male** | ***Celsr3^+/+^/ Celsr3^S1894Rfs*2/+^*** **Female** | ***Celsr3^+/+^/ Celsr3^S1894Rfs*2/+^*** **Male** | ***Wwc1^+/+^/ Wwc1^W88C^*** **Female** | ***Wwc1^+/+^/ Wwc1^W88C/+^*** **Male** |
| **Prepulse inhibition (PPI) of the acoustic startle reflex** | | | | | | | |
| **PPI (%)** | **Two-way ANOVA** | ***n=14/11*** | ***n=16/11*** | ***n=13/25*** | ***n=11/16*** | ***n=17/20*** | ***n=13/29*** |
| | *Genotype main effect* | *F (1, 75)=16.95, p=0.0005* | *F(1, 69)=13.46, p<0.0001* | *F(1, 108)=8.34, p=0.0047* | *F(1, 75)=8.7, p=0.0042* | *F(1, 105)=7.69, p=0.0066* | *F(1, 120)=0.818, p=0.367* |
| **Startle amplitude** | **Unpaired t-test** | *p=0.798* | *p=0.403* | *p=0.038* | *p=0.146* | *p=0.96* | *p=0.91* |

| | | | | | | | |
|---|---|---|---|---|---|---|---|
| **PPI (%) Pretreatment with Aripiprazole** | **Multiple Linear Regression (Factor III)** | ***Vehicle treated: n=16/11*** <br> ***Drug treated: n=13/13*** | | ***Vehicle treated: n=18/16*** <br> ***Drug treated: n=13/13*** | | | |
| | *Model* | *F(11, 147)=6.49, p<0.00001, $R^2$=0.28* | | F(11, 168)=6.987, *p<0.00001, $R^2$=0.27* | | | |
| | *Genotype* | *t=* -1.69, *p=0.093* | | *t=* -1.868, *p=0.064* | | | |
| | *Treatment* | *t*=0.776, *p=0.44* | | *t*=0.463, *p=0.66* | | | |
| | *Treatment x Genotype* | *t*=2.419, *p=0.017* | | *t*=2.507, *p=0.013* | | | |
| | ***Tukey's Multiple Comparisons test*** | **Vehicle only:** <br> ***$Celsr3^{+/+}$ vs. $Celsr3^{C1906Y/+}$*** <br> *p=0.02* <br><br> **Vehicle vs. Drug:** <br> ***$Celsr3^{+/+}$ vs. $Celsr3^{C1906Y/+}$*** <br> *p=0.34* <br><br> **Vehicle vs. Drug:** <br> ***$Celsr3^{C1906Y/+}$ vs. $Celsr3^{C1906Y/+}$*** <br> *p=0.14* <br><br> **Vehicle vs. Drug:** <br> ***$Celsr3^{+/+}$ vs. $Celsr3^{+/+}$:*** <br> *p=0.61* | | **Vehicle only:** <br> ***$Celsr3^{+/+}$ vs. $Celsr3^{S1894Rfs*2/+}$*** <br> *p=0.006* <br><br> **Vehicle vs. Drug:** <br> ***$Celsr3^{+/+}$ vs. $Celsr3^{S1894Rfs*2/+}$*** <br> *p=0.74* <br><br> **Vehicle vs. Drug:** <br> ***$Celsr3^{C1906Y/+}$ vs. $Celsr3^{C1906Y/+}$*** <br> *p=0.17* <br><br> **Vehicle vs. Drug:** <br> ***$Celsr3^{+/+}$ vs. $Celsr3^{+/+:}$*** <br> *p=0.46* | | | |
| **Rotarod** | | | | | | | |
| **Latency to fall** | **RM Two-way ANOVA** | ***n=20/14*** | ***n=17/15*** | ***n=20/18*** | ***n=17/26*** | ***n=27/29*** | ***n=19/40*** |
| | Genotype main effect | *F(1, 32)=3.9, p=0.056* | *F(1, 30)=6.6, p=0.015* | *F(1, 36)=0.11, p=0.75* | *F(1, 41)=4.4 p=0.051* | *F(1, 55)=0.21, p=0.66* | *F(1, 57)=0.03, p=0.86* |
| **Open field – locomotion** | | | | | | | |

| | | | | | | | |
|---|---|---|---|---|---|---|---|
| **Total distance travelled (summed at each time point)** | **RM Two-way ANOVA** | ***n=16/16*** | ***n=15/17*** | ***n=18/25*** | ***n=12/29*** | ***n=12/21*** | ***n=13/18*** |
| | *Time x Genotype* | *F(2, 60)=6.03, p=0.004* | *F(2, 60)=2.85, p=0.066* | *F(2,82)=0.78, p=0.46* | *F(2,78)=0.55, p=0.58* | *F(2,64)=0.18, p=0.83* | *F(2, 58)=5.5, p=0.006* |
| | *Genotype main effect* | *F(1, 30)=6.11, p=0.019* | *F(1, 30)=2.814, p=0.104* | *F(1,41)=2.8, p=0.102* | *F(1,39)=0.61, p=0.44* | *F(1,32)=0.02, p=0.89* | *F(1,29)=4.2, p=0.049* |
| | ***Bonferroni's Multiple Comparisons Test*** | ***10**, p=0.55*<br>***20**, p=0.07*<br>***30**, p=0.004* | ***0**, p>0.99*<br>***20**, p=0.36*<br>***30**, p=0.07* | ***0**, p=0.77*<br>***20**, p=0.25*<br>***30**, p=0.21* | ***10**, p>0.99*<br>***20**, p>0.99*<br>***30**, p=0.85* | ***10**, p>0.99*<br>***20**, p>0.99*<br>***30**, p>0.99* | ***10**, p=0.46*<br>***20**, p=0.19*<br>***30**, p=0.10* |
| **Latency to enter center** | **Unpaired t-test** | *p=0.61* | *p=0.59* | *p=0.073* | *p=0.67* | *p=0.66* | *p=0.74* |
| **Total Central zone entries (summed at each time point)** | **RM Two-way ANOVA** | | | | | | |
| | *Time x Genotype* | F(2, 60)=5.69, *p=0.005* | F(2, 60)=2.04, *p=0.139* | F(2, 82)=2.2 *p=0.12* | F(2, 78)=0.35, *p=0.965* | F(2, 64)=1.6, p=0.21 | F(2, 58)=0.056, *p=0.95* |
| | *Genotype main effect* | F(1, 30)=1.88, *p=0.18* | F(1, 30)=1.3, *p=0.266* | *F(1, 41)=3.6, p=0.065* | F(1,39)=0.1922, *p=0.651* | F(1, 32)=0.15, *p=0.703* | F(1, 29)=1.1, *p=0.29* |
| | ***Bonferroni's Multiple Comparisons Test*** | *10, p=0.85*<br>***20**, p=0.25*<br>***30**, p=0.09* | ***10**, p=0.99*<br>***20** p=0.66*<br>***30**, p=0.37* | ***10**, p=0.62*<br>***20**, p=0.26*<br>***30**, p=0.052* | ***0**, p>0.99*<br>***20**, p>0.99*<br>***30**, p>0.99* | ***10**, p>0.99*<br>***20**, p>0.99*<br>***30**, p>0.99* | ***10**, p=0.27*<br>***20**, p=0.85*<br>***30**, p>0.99* |
| | | | | **Open Field Rearing** | | | |
| **Total Rearing Events (summed at each time point)** | **RM Two-way ANOVA** | | | | | | |
| | Time x Genotype | *F(2, 60)=5.99, p=0.004* | *F(2, 60)=0.72, p=0.49* | *F(2, 82)=5.8, p=0.004* | *F(2, 78)=3.6, p=0.03* | *F(2, 64)=0.02, p=0.98* | *F(2, 58)=2.2, p=0.12* |
| | *Genotype main effect* | *F(1, 30)=10.77, p=0.003* | *F(1, 30)=7.44, p=0.011* | *F(1, 41)=10.8, p=0.002* | *F(1, 39)=4.6, p=0.038* | *F(1, 32)=0.35, p=0.56* | *F(1, 29)=4.2, p=0.048* |
| | **Bonferroni's Multiple Comparisons Test** | ***10**, p=0.24*<br>***20**, p=0.01*<br>***30**, p=0.002* | ***10**, p=0.24*<br>***20**, p=0.028*<br>***30**, p=0.026* | ***10**, p=0.25*<br>***20**, p=0.008*<br>***30**, p=0.0002* | ***10**, p>0.99*<br>***20**, p=0.13*<br>***30**, p=0.013* | ***10**, p>0.99*<br>***20**, p>0.99*<br>***30**, p>0.99* | ***10**, p=0.18*<br>***20**, p=0.24*<br>***30**, p=0.24* |

| | | | | | | | |
|---|---|---|---|---|---|---|---|
| **Rearing events across binned intervals** | **RM Two-way ANOVA**<br>*Genotype main effect* | *F(1,30)=9.1, p=0.005* | *F(1,30)=3.6, p=0.067* | *F(1, 41)=9.5, p=0.0035* | *F(1,39)=5.8, p=0.021* | *F(1,32)=0.026, p=0.87* | *F(1, 29)=3.2, p=0.066* |
| | **Bonferroni's multiple comparisons test** | ***0-10**, p=0.002*<br>***10-20**, p=0.046*<br>***20-30**, p=0.074* | ***0-10**, p=0.005*<br>***10-20**, p=0.16*<br>***20-30**, p=0.95* | ***0-10**, p=0.004*<br>***10-20**, p=0.043*<br>***20-30**, p=0.098* | ***0-10**, p=0.19*<br>***10-20**, p=0.066*<br>***20-30**, p=0.11* | ***0-10**, p>0.99*<br>***10-20**, p>0.99*<br>***20-30**, p>0.99* | ***0-10**, p=0.14*<br>***10-20**, p=0.36*<br>***20-30**, p=0.42* |
| **Total time rearing (summed at each time point)** | **RM Two-way ANOVA** | | | | | | |
| | *Time x genotype* | *F(2, 60)=2.31, p=0.11* | *F(2, 60)=3.9, p=0.024* | *F(2, 82)=12.19, p<0.0001* | *F(2, 78)=2.9, p=0.059* | *F(2, 64)=0.09, p=0.9* | *F(2, 58)=12.5, p=0.083* |
| | *Genotype main effect* | *F(1, 30)=4.3, p=0.045* | *F(1, 30)=0.44, p=0.52* | *F(1, 41)=8.09 p=0.0069* | *F(1, 39)=4.9, p=0.03* | *F(1, 32)=0.0001, p=0.97* | *F(1, 29)=3.6, p=0.065* |
| | **Bonferroni's Multiple Comparisons test** | ***10**, p=0.85*<br>***20**, p=0.25*<br>***30**, p=0.02* | ***10**, p=0.21*<br>***20**, p=0.74*<br>***30**, p>0.99* | ***10**, p=0.99*<br>***20**, p=0.203*<br>***30**, p<0.0001* | ***10**, p>0.99*<br>***20**, p=0.13*<br>***30**, p=0.012* | ***10**, p>0.99*<br>***20**, p>0.99*<br>***30**, p>0.99* | ***10**, p>0.99*<br>***20**, p=0.38*<br>***30**, p=0.025* |
| **Rearing time across binned intervals** | **RM Two-way ANOVA** | | | | | | |
| | *Genotype main effect* | *F(1, 30)=4.0, p=0.054* | *F(1,30)=0.45, p=0.51* | *[F(1, 41)=11.43, p=0.002* | *F(1, 39)=6.1, p=0.018* | *[F(1, 32)=0.18, p=0.89* | *F(1, 29)=1.8, p=0.076* |
| | **Bonferroni's Multiple Comparisons Test** | ***0-10**, p=0.0.045*<br>***10-20**, p=0.21*<br>***20-30**, p=0.73* | ***0-10**, p=0.20*<br>***10-20**, p=0.74*<br>***20-30**, p>0.99* | ***0-10**, p=0.13*<br>***10-20**, p=0.22*<br>***20-30**, p=0.0005* | ***0-10**, p=0.17*<br>***10-20**, p=0.04*<br>***20-30**, p=0.39* | ***0-10**, p>0.99*<br>***10-20**, p>0.99*<br>***20-30**, p>0.99* | ***0-10**, p=0.55*<br>***10-20**, p=0.49*<br>***20-30**, p=0.13* |
| **Cumulative Perimeter rearing events** | **Mann Whitney test** | *p=0.001* | *p=0.023* | *p=0.08* | *p=0.06* | *p=0.45* | *p=0.12* |
| **Cumulative Center rearing events** | **Mann Whitney test** | *p=0.25* | *p=0.85* | *p=0.08* | *p=0.24* | *p=0.34* | *p=0.24* |

| | | | | |
|---|---|---|---|---|
| **Rearing events across binned intervals**<br>**Aripiprazole pretreatment** | **Multiple Linear Regression (Factor III)** | **Vehicle treated:**<br>***n=12/10***<br>**Drug treated:**<br>***n=10/10*** | | **Vehicle treated:**<br>***n=12/11***<br>**Drug treated:**<br>***n=10/10*** |
| | *Model* | $F(11, 114)=17.99$,<br>$p<0.00001$,<br>$R^2=0.59$ | | [$F(11, 117)=18.13$,<br>$p<0.00001$,<br>$R^2=0.59$ |
| **Rearing events** | *Genotype* | $t=7.41$, $p<0.0001$ | | $t=6.57$, $p<0.0001$ |
| | *Treatment* | $t=11.87$, $p<0.0001$ | | $t=11.88$, $p<0.0001$ |
| | *Genotype x Treatment interaction* | $t=1.06$, $p=0.291$ | | $t=1.11$, $p=0.269$ |
| | **Tukey's Multiple Comparisons Test** | **Vehicle only:**<br>***Celsr3$^{+/+}$ vs. Celsr3$^{C1906Y/+}$***<br>*0-10, p=0.11*<br>*10-20, p=0.007*<br>*20-30, p=0.02*<br><br>**Vehicle vs. Drug:**<br>***Celsr3$^{+/+}$ vs. Celsr3$^{C1906Y/+}$***<br>*0-10, p=0.99*<br>*10-20, p=0.58*<br>*20-30, p=0.51*<br><br>**Vehicle vs. Drug:**<br>***Celsr3$^{C1906Y/+}$ vs. Celsr3$^{C1906Y/+}$***<br>*0-10, p=0.006*<br>*10-20, p<0.0001*<br>*20-30, p<0.0001*<br><br>**Vehicle vs. Drug:**<br>***Celsr3$^{+/+}$ vs. Celsr3$^{+/+}$*:**<br>*0-10, p=0.0024*<br>*10-20, p=0.0033*<br>*20-30, p<0.0001* | | **Vehicle only:**<br>***Celsr3$^{+/+}$ vs. Celsr3$^{S1894Rfs*2/+}$***<br>*0-10, p=0.03*<br>*10-20, p=0.14*<br>*20-30, p=0.12*<br><br>**Vehicle vs. Drug:**<br>***Celsr3$^{+/+}$ vs. Celsr3$^{S1894Rfs*2/+}$***<br>*0-10, p=0.99*<br>*10-20, p=0.79*<br>*20-30, p=0.47*<br><br>**Vehicle vs. Drug:**<br>***Celsr3$^{S1894Rfs*2/+}$vs. Celsr3$^{S1894Rfs*2/+}$***<br>*0-10, p<0.0001*<br>*10-20, p=0.05*<br>*20-30, p=0.01*<br><br>**Vehicle vs. Drug:**<br>***Celsr3$^{+/+}$ vs. Celsr3$^{++}$*:**<br>*0-10, p=0.015*<br>*10-20, p=0.003*<br>*20-30, p=0.0001* |

| | | | | | | | |
|---|---|---|---|---|---|---|---|
| **Open field - grooming** | | | | | | | |
| **Cumulative Grooming events** | **Mann Whitney test** | ***n=13/12***<br><br>*p=0.001* | ***n=10/18***<br><br>*p=0.08* | ***n=14/21***<br><br>*p=0.0005* | ***n=12/17***<br><br>*p=0.045* | ***n=11/16***<br><br>*p=0.518* | ***n=11/29***<br><br>*p=0.704* |
| **Cumulative Time spent grooming** | | *p=0.05* | *p=0.87* | *p=0.001* | *p=0.62* | *p=0.34* | *p=0.41* |
| **Mean time per grooming events** | | *p=0.65* | *p=0.25* | *p=0.61* | *p=0.59* | *p=0.11* | *p=0.36* |
| **Marble burying assay** | | | | | | | |
| **Cumulative marbles buried** | **Mann Whitney test** | ***n=19/13***<br><br>*p=0.032* | ***n=13/13***<br><br>*p=0.6* | ***n=14/16***<br><br>*p=0.07* | ***n=9/23***<br><br>*p=0.39* | ***n=15/21***<br><br>*p=0.38* | ***n=15/17***<br><br>*p=0.44* |
| ***Instrumental learning task – fixed ratio reinforcement*** | | | | | | | |
| **Rate of rewards earned** | **Regression Analysis**<br><br>*Linear regression, Day 1*<br><br>*Non-linear regression, Day 2-4* | | ***n=26/30***<br><br>*p<0.0001*<br><br>*p<0.0001* | ***n=27/22***<br><br>*p<0.0001*<br><br>*p<0.0001* | | | |
| **Mean time to obtain 30 rewards** | **RM Two-way ANOVA**<br><br>Genotype main effect | | *F(1, 219)=4.998, p=0.026* | *F(1, 191)=11.43, p=0.0009* | | | |
| **Latency to retrieve first reward (Day 2-4)** | **ANOVA with ordinary least squares method for linear regression** | | *p=0.093* | *p=0.075* | | | |

**Table S1:** Statistical analyses and *p*-values for prepulse inhibition, open field arena, and conditioned fixed reinforcement by sex and genotype. Red and blue indicate an increase or decrease, respectively, in comparison to control.

### Table S2 – Annotations and statistics for Motion Sequencing syllable usage.

| | | | |
|---|---|---|---|
| | $p < 0.05$ (up in mutant) | | $p < 0.05$ (down in mutant) |
| | $0.05 < p < 0.1$ (up in mutant) | | $0.05 < p < 0.1$ (down in mutant) |

| | | Test group | | | | | |
|---|---|---|---|---|---|---|---|
| | | ***Celsr3***$^{C1906Y}$ **F** | ***Celsr3***$^{C1906Y}$ **M** | ***Celsr3***$^{p.S1894Rfs*2}$ **F** | ***Celsr3***$^{p.S1894Rfs*2}$ **M** | ***Wwc1***$^{W88C}$ **F** | ***Wwc1***$^{W88C}$ **M** |
| **Syllable ID#** | **Annotation** | WT n=26<br>MT n=22 | WT n=21<br>MT n=20 | WT n=25<br>MT n=33 | WT n=28<br>MT n=48 | WT n=11<br>MT n=16 | WT n=12<br>MT n=26 |
| **rearing syllables** | | | | | | | |
| 1 | low rear • | ns | 0.0721 | ns | ns | ns | ns |
| 3 | rear down, head turn | ns | ns | ns | 0.0706 | ns | ns |
| 5 | step forward, low rear • | ns | ns | 0.0627 | 0.0275 | ns | ns |
| 6 | come down from wall rear • | 0.0181 | 0.0240 | ns | 0.0017 | ns | ns |
| 8 | down from low rear | ns | ns | ns | ns | ns | ns |
| 10 | down from low rear | 0.0814 | ns | ns | ns | ns | ns |
| 17 | wall assisted rear • | 0.0009 | 0.0037 | ns | 0.0640 | ns | ns |
| 25 | pre rear stance | ns | ns | ns | ns | ns | ns |
| 26 | unassisted high rear | ns | ns | ns | ns | ns | ns |
| 28 | nose up in rear | ns | ns | ns | ns | ns | ns |
| 29 | nose up in low rear | ns | 0.0751 | ns | ns | ns | ns |
| 30 | wall assisted rear | ns | ns | ns | ns | ns | ns |
| 31 | unassisted rear | ns | ns | ns | 0.0590 | ns | ns |
| 32 | unassisted rear | ns | ns | ns | ns | ns | ns |
| 37 | wall assisted rear • | ns | ns | 0.1000 | 0.0056 | ns | ns |
| 40 | unassisted rear | ns | ns | ns | ns | ns | ns |
| 41 | wall assisted rear | ns | ns | ns | ns | ns | ns |
| 42 | wall assisted rear • | ns | ns | 0.0354 | ns | ns | ns |
| 44 | wall assisted rear | ns | ns | ns | ns | ns | ns |
| 45 | wall assisted high rear • | ns | 0.0324 | ns | ns | ns | ns |
| 47 | wall assisted rear • | 0.0372 | 0.0469 | ns | ns | ns | ns |
| 49 | wall assisted rear • | 0.0610 | ns | 0.0404 | ns | ns | ns |
| 50 | wall assisted rear | ns | ns | 0.0500 | ns | ns | ns |
| 51 | unassisted low rear | ns | ns | ns | ns | ns | ns |
| 53 | unassisted high rear | ns | ns | ns | ns | ns | ns |
| 59 | unassisted high rear | ns | ns | ns | ns | ns | ns |
| 66 | low rear | ns | 0.0548 | ns | ns | ns | ns |
| 67 | wall assisted rear | ns | ns | ns | ns | ns | ns |
| **grooming syllables** | | | | | | | |
| 23 | groom, on hind legs | 0.0934 | ns | ns | ns | ns | ns |
| 60 | groom • | ns | ns | 0.0358 | ns | ns | ns |

| Syllable ID# | Annotation | *Celsr3*$^{C1906Y}$ F | *Celsr3*$^{C1906Y}$ M | *Celsr3*$^{p.S1894Rfs*2}$ F | *Celsr3*$^{p.S1894Rfs*2}$ M | *Wwc1*$^{W88C}$ F | *Wwc1*$^{W88C}$ M |
|---|---|---|---|---|---|---|---|
| **grooming syllables cont'd** | | | | | | | |
| 62 | groom face | ns | ns | ns | ns | ns | ns |
| 63 | groom | ns | ns | 0.0587 | ns | ns | ns |
| 64 | groom face | ns | ns | 0.0937 | ns | ns | ns |
| 65 | groom | ns | ns | 0.0937 | ns | ns | 0.0543 |
| **locomotion syllables** | | | | | | | |
| 4 | locomotion forward • | 0.0903 | ns | 0.0587 | 0.0064 | ns | ns |
| 9 | locomotion forward • | 0.0151 | ns | ns | 0.0213 | ns | 0.0694 |
| 11 | locomotion forward • | ns | ns | 0.0186 | 0.0980 | ns | ns |
| 12 | locomotion forward • | 0.0461 | ns | ns | ns | ns | ns |
| 16 | locomotion forward, head up | ns | ns | ns | ns | ns | ns |
| 18 | locomotion forward • | 0.0417 | 0.0993 | ns | ns | ns | ns |
| 19 | locomotion forward | ns | ns | ns | ns | ns | ns |
| 27 | forward locomotion | ns | ns | ns | ns | ns | ns |
| 34 | locomotion forward | ns | ns | ns | ns | ns | ns |
| 35 | locomotion forward | 0.0517 | ns | ns | ns | ns | ns |
| 36 | locomotion forward | ns | 0.0657 | ns | ns | ns | ns |
| 38 | locomotion forward | ns | ns | ns | ns | ns | ns |
| 39 | forward locomotion, head up | ns | ns | ns | ns | ns | ns |
| 54 | locomotion around edge | ns | ns | 0.0531 | ns | ns | ns |
| 58 | locomotion around edge | ns | ns | 0.0657 | ns | ns | ns |
| **pause syllables** | | | | | | | |
| 14 | pause • | 0.0061 | 0.0227 | 0.0303 | 0.0013 | ns | ns |
| 20 | pause, head down | 0.0627 | 0.0721 | ns | ns | ns | ns |
| 48 | pause • | 0.0159 | 0.0192 | ns | 0.0355 | ns | ns |
| 52 | pause • | ns | 0.0163 | 0.0320 | 0.0994 | ns | ns |
| 55 | pause • | 0.0047 | 0.0202 | 0.0123 | 0.0009 | ns | ns |
| 56 | pause • | 0.0496 | 0.0469 | 0.0044 | 0.0084 | ns | ns |
| 57 | pause • | ns | ns | 0.0420 | 0.0980 | ns | ns |
| 68 | pause in center | ns | ns | ns | ns | ns | ns |
| 70 | pause | - | - | - | ns | - | - |
| **other syllables** | | | | | | | |
| 0 | scrunch | ns | 0.0845 | 0.1000 | ns | ns | ns |
| 2 | scrunch, head down | ns | ns | ns | ns | ns | ns |
| 7 | scrunch • | ns | ns | ns | 0.0040 | ns | ns |
| 13 | left turn | ns | ns | ns | 0.0889 | ns | ns |
| 15 | head down | ns | ns | ns | ns | ns | ns |
| 21 | scrunch | ns | 0.0960 | 0.1000 | ns | ns | ns |
| 22 | step forward, head up • | 0.0446 | ns | ns | ns | ns | ns |

| Syllable ID# | Annotation | *Celsr3*$^{C1906Y}$ F | *Celsr3*$^{C1906Y}$ M | *Celsr3*$^{p.S1894Rfs*2}$ F | *Celsr3*$^{p.S1894Rfs*2}$ M | *Wwc1*$^{W88C}$ F | *Wwc1*$^{W88C}$ M |
|---|---|---|---|---|---|---|---|
| **other syllables cont'd** | | | | | | | |
| 24 | head bob ● | 0.0934 | ns | 0.0434 | 0.0650 | ns | ns |
| 33 | head bob | ns | ns | ns | ns | ns | ns |
| 43 | scrunch, head down | ns | ns | ns | ns | ns | ns |
| 46 | head bob ● | 0.0010 | 0.0429 | 0.0057 | 0.0160 | ns | ns |
| 61 | jump ● | ns | ns | 0.0041 | ns | ns | ns |
| 69 | jump at side ● | ns | ns | 0.0088 | ns | ns | ns |

**Table S2: Ethological classification of significantly changed MoSeq syllables.** Table of significantly changed syllables across the different sexes and mutations studied. Column “syllable” refers to the usage-sorted syllable ID, column “description” is a human generated behavioral annotation, column “class” is a high-level behavioral class assigned (e.g., locomotion, pause, rear, groom, jump). Each column for a given mutation and sex illustrates syllables which are significantly changed in the mutant relative to respective control. The color and text within individual table cells indicate the direction of change in the mutant relative to the respective control (red/up indicates up-regulation in the mutant, blue/down indicates down-regulation in the mutant).
